# Mapping snoRNA-target RNA interactions in an RNA binding protein-dependent manner with chimeric eCLIP

**DOI:** 10.1101/2024.09.19.613955

**Authors:** Zhuoyi Song, Bongmin Bae, Simon Schnabl, Fei Yuan, Thareendra De Zoysa, Maureen Akinyi, Charlotte Le Roux, Karine Choquet, Amanda Whipple, Eric Van Nostrand

**Author notes:** Authors contributed equally (Zhuoyi Song, Bongmin Bae).

## Abstract

Small nucleolar RNAs (snoRNAs) are non-coding RNAs that function in ribosome and spliceosome biogenesis, primarily by guiding modifying enzymes to specific sites on ribosomal RNA (rRNA) and spliceosomal RNA (snRNA). However, many orphan snoRNAs remain uncharacterized, with unidentified or unvalidated targets, and studies on additional snoRNA-associated proteins are limited. We adapted an enhanced chimeric eCLIP approach to comprehensively profile snoRNA-target RNA interactions using both core and accessory snoRNA binding proteins as baits. Using core snoRNA binding proteins, we confirmed most annotated snoRNA-rRNA and snoRNA-snRNA interactions in mouse and human cell lines and called novel, high-confidence interactions for orphan snoRNAs. While some of these interactions result in chemical modification, others may have modification-independent functions. We then showed that snoRNA ribonucleoprotein complexes containing certain accessory proteins, like WDR43 and NOLC1, enriched for specific subsets of snoRNA-target RNA interactions with distinct roles in ribosome and spliceosome biogenesis. Notably, we discovered that SNORD89 guides 2’-O-methylation at two neighboring sites in U2 snRNA that are important for activating splicing, but also appear to ensure imperfect splicing for a subset of near-constitutive exons. Thus, chimeric eCLIP of snoRNA-associating proteins enables a comprehensive framework for studying snoRNA-target interactions in an RNA binding protein-dependent manner, revealing novel interactions and regulatory roles in RNA biogenesis.

## Introduction

Small nucleolar RNAs (snoRNAs) constitute a class of non-coding RNAs primarily known for their fundamental roles in ensuring the proper biogenesis of ribosomal RNA (rRNA). SnoRNAs can be categorized into box C/D and box H/ACA snoRNAs, distinguished by the presence of conserved box motif sequences and structural features. In addition, a class of snoRNAs known as small Cajal body-specific snoRNAs (scaRNAs), which contain either C/D, H/ACA, or hybrid (both) motifs, are involved in the biogenesis of spliceosomal RNAs (snRNAs). During their maturation, snoRNAs associate with core RNA-binding proteins (snoRBPs) to form functional snoRNPs. Specifically, C/D snoRNAs associate with FBL, NOP56, NOP58, and SNU13 (NHP2L1/15.5K), while H/ACA snoRNAs associate with DKC1, NOP10, NHP2 and GAR1 (Fig. 1A). SnoRNPs translocate to the nucleolus or Cajal body, where they interact with nascent pre-rRNA or snRNA [1]. The antisense elements of snoRNAs engage in base pairing with their RNA targets, orchestrating the precise positioning of 2’-O-methylation (Nm) by C/D snoRNPs or isomerization of uridine to pseudouridine (Ψ) by H/ACA snoRNPs. Human rRNA contains more than 200 nucleotides that undergo snoRNA-guided chemical modification [2]. Some of these chemical modifications cluster at functionally important regions of the ribosome, such as the peptidyl transferase center, tRNA binding sites, and the interface between the small and large subunits, and they contribute to the stabilization of rRNA folding, facilitation of efficient ribosome assembly, export of ribosomal subunits, and binding interactions with translation factors [3–5].

**Figure 1.**
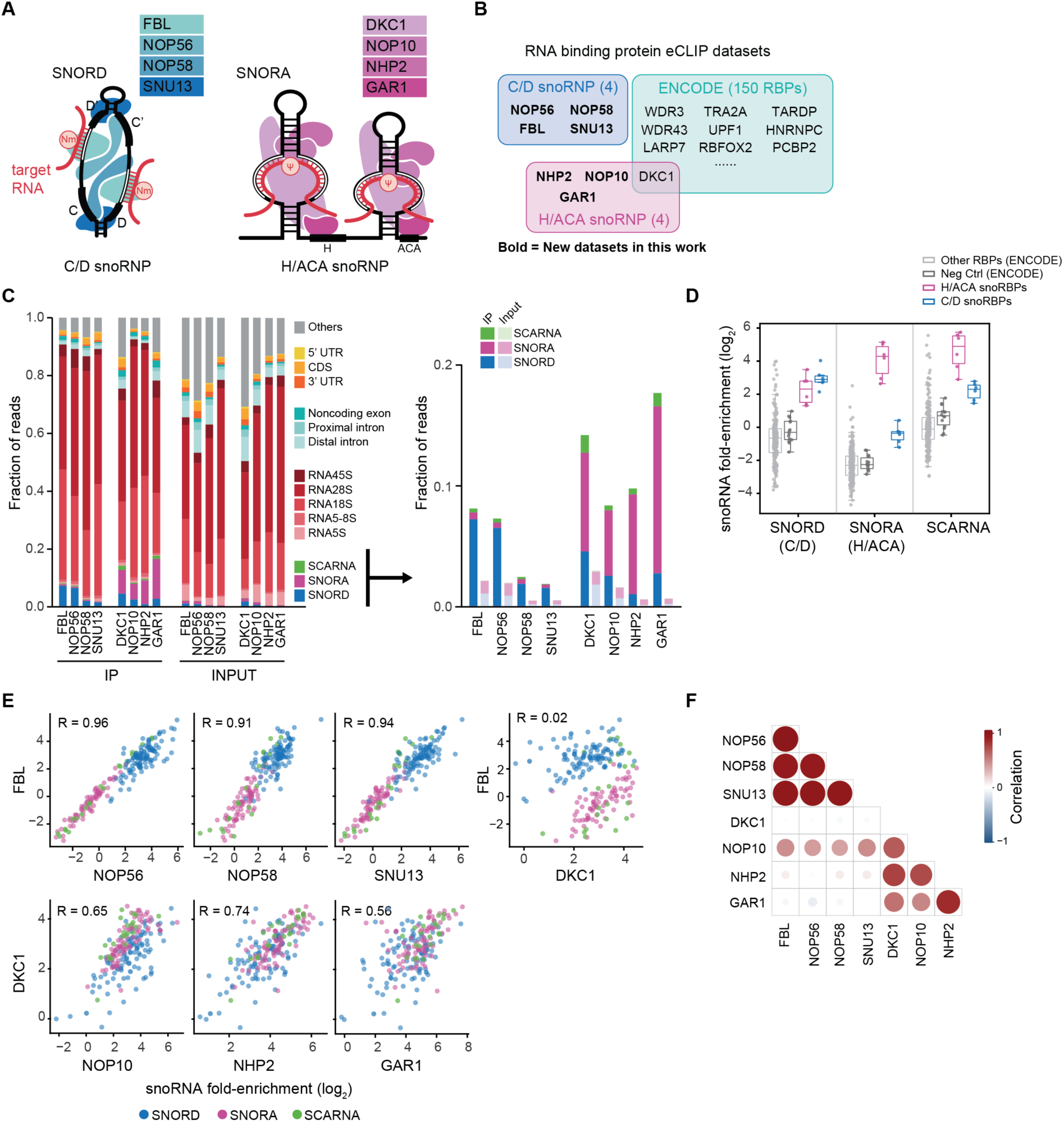
Core snoRNP proteins show consistent RNA interactomes. **(A)** Schematic of (left) C/D and (right) H/ACA snoRNA-protein complexes. Red line represents the target RNA complementary to the snoRNA antisense element. **(B)** Overview of snoRNA binding proteins with RNA interactomes profiled by ENCODE or (boldface) this work. **(C)** Distribution of reads to (left) major RNA classes and (right) snoRNA classes captured by eCLIP in K562 cells of the indicated core snoRBPs. **(D)** Fold-enrichment of different classes of snoRNAs in eCLIP datasets (both from ENCODE and generated in this study). Colors indicate different groups of protein baits (control - FLAG or V5). **(E)** Correlation of snoRNA fold-enrichment across (top) core C/D snoRBPs or (bottom) core H/ACA snoRBPs from eCLIP. Pearson correlations (R) are indicated. (top right) Correlation between core C/D and H/ACA snoRBPs shown for contrast. Datapoints with less than 5 reads in IP were discarded. **(F)** Bubble plot showing pairwise correlation in snoRNA fold-enrichment between snoRBPs. Circle size and color indicate value of Pearson correlation coefficient (R).

Over the years, research efforts have successfully identified snoRNA-target pairs for most known 2’-O-methylation and pseudouridine sites in rRNA. Advances in mass spectrometry approaches and high-throughput assays have facilitated the discernment of nucleotides subject to chemical modifications [2]. Additionally, bioinformatics tools have been employed to associate numerous snoRNAs with specific rRNA modification sites [6–9] . The development of RNA-RNA interaction mapping tools have validated additional snoRNA-rRNA interactions [10–15]. However, despite these collective efforts, the targets for a considerable subset of snoRNAs, known as orphan snoRNAs, have yet to be identified or experimentally validated.

Beyond their role in rRNA modifications, C/D snoRNAs can guide Nm in other RNA targets. This extends to snRNAs, tRNAs, and even a few mRNAs [13,16–20]. Furthermore, C/D snoRNAs can act in a chaperone-like fashion independent of chemical modification. For instance, base pairing interactions between snoRNAs and pre-rRNA contribute to rRNA processing and assembly by influencing pre-rRNA conformation [21,22]. Additionally, interactions between snoRNAs and pre-mRNA can contribute to splicing and 3’ end processing [20,23,24], further highlighting the multifaceted roles of these snoRNAs beyond guiding chemical modifications. In many instances, these non-traditional functions require the assembly of additional proteins to snoRNA complexes [24–26]; however, the landscape of RNA-binding proteins (RBPs) and their snoRNA-interacting networks beyond the core snoRBP complexes remain undercharacterized.

Here we performed a deep, large-scale profiling of RNA interactions for all eight core RNA-binding protein components of the C/D and H/ACA snoRNA complexes. We then adopted chimeric eCLIP to directly capture snoRNA-target RNA interactions, utilizing both established core proteins and lesser-known snoRNA-associated RBPs as bait. Our results demonstrate that chimeric eCLIP successfully corroborated canonical snoRNA interactions with rRNAs and snRNAs. Moreover, we discovered novel interactions of orphan snoRNAs with rRNA and snRNA, a subset of which mediate chemical modification at the target site. Using non-core snoRNA-associated RBPs as baits in chimeric eCLIP revealed specific subnetworks of snoRNAs, suggesting specific roles of snoRNAs in distinct stages of snRNA or rRNA biogenesis. We further show that one such snoRNA, SNORD89, plays a key role in 2’-O-methylation of the G11 and G12 sites in U2 snRNA, which appears to be essential for maintaining imperfect splicing. Together, this study presents a systematic approach to investigate snoRNA-target interactions, revealing novel roles in RNA biogenesis and splicing regulation, and highlighting the potential for further discoveries in snoRNA biology.

## Results

### Core snoRNP proteins show consistent RNA interactomes

Previous profiling of the RNA interactomes of core C/D snoRNP proteins FBL, NOP56, and NOP58 by CLIP-seq led to the characterization of novel snoRNAs as well as the identification of candidate interactions for orphan snoRNAs [27,28]. However, of the four core H/ACA snoRNP proteins, the RNA interactome has only been profiled for DKC1 [27,29]. To expand upon this prior work, we set out to use the updated eCLIP framework that allows deeper recovery of unique RNA molecules as well as quantitative normalization against input controls to comprehensively profile the direct interactions for the C/D and H/ACA snoRNP complex members [30]. We obtained and performed IP-western blot validation of antibodies against the four core proteins for both C/D (FBL, NOP56, NOP58, and SNU13) and H/ACA (DKC1, NOP10, NHP2, and GAR1) snoRNP complexes respectively (Fig. S1A), followed by eCLIP in K562 cells to facilitate contrast analysis with other ENCODE RBP datasets (Fig. 1B).

Basic analysis of snoRBP eCLIP indicated successful enrichment of snoRNAs (Fig. 1C,D). Similar to previous observations with PAR-CLIP [27], eCLIP for FBL, NOP56, NOP58, and SNU13 each showed significant enrichment for C/D snoRNAs versus paired input (≥ 6.5-fold) (Fig. 1C). Over 5% of reads mapped to C/D snoRNAs for FBL and NOP56, with an additional 80% of reads mapping to either precursor or mature rRNA; NOP58 and SNU13 had significantly enriched but lower frequency of reads mapping to C/D snoRNAs (>1.5%) (Fig. 1C). The C/D snoRBPs showed an average 1.5-fold depletion for H/ACA snoRNAs compared to size-matched inputs, suggesting that the C/D complex shows specificity for binding to C/D snoRNAs (Fig. 1C,D). Similarly, all four H/ACA snoRBPs showed significant enrichment for H/ACA snoRNAs (≥ 6.2-fold) (Fig. 1C,D). Over 5% of reads mapped to H/ACA snoRNAs for each RBP and another 71% to rRNA (Fig. 1C). Surprisingly, all four H/ACA snoRBPs also showed enrichment for C/D snoRNAs (≥ 2.5-fold) (Fig. 1C,D), which may indicate a broader role for H/ACA snoRBPs or reflect low-level co-immunoprecipitation of FBL in these experiments (Fig. S1B).

Previous research has observed examples of sub-complex specificity, as SNU13 is involved in other RNA-protein complexes such as the U4/U6.U5 tri-snRNP [31] in addition to its role in snoRNP assembly. Consistent with this, eCLIP of SNU13 but not the other C/D snoRBPs showed enrichment at the 5’ stem loop of the U4 snRNA similar to other U4-interacting tri-snRNP factors (Fig. S1C). However, when we performed transcriptome-wide analysis comparing the enrichment of individual snoRNAs, we observed striking pair-wise correlation in per-snoRNA enrichment among all core C/D snoRBPs (all Pearson R > 0.90) and significant correlation in enrichment among all core H/ACA snoRBPs (R > 0.50) (Fig. 1E,F), which was independent of FBL or DKC1 co-immunoprecipitation (Fig. S1B). In contrast, there is little correlation in per-snoRNA enrichment between C/D and H/ACA snoRBPs, such as FBL and DKC1 (R = 0.02) (Fig. 1E,F). To consider these results in the context of outgroup (non-snoRNA-related) RBPs, we utilized available ENCODE eCLIP data to compare the correlation of enrichment for individual snoRNAs across 149 other RBPs. Although the majority of RBP and negative control (using FLAG or V5 pulldown in wildtype cells) datasets were depleted for snoRNAs (Fig. 1D), we surprisingly observed a significant per-snoRNA correlation with FBL across RBP datasets (median R = 0.62) and negative controls (median R = 0.76) that may reflect low-level co-precipitation of the C/D snoRNP complex across eCLIP experiments (Fig. S1B,D). However, the correlation between pairs of core C/D or H/ACA snoRBPs was higher than with nearly any other RBPs (Fig. S1D), providing further evidence that the four core C/D and four core H/ACA snoRBPs coordinately associate with C/D and H/ACA snoRNAs, respectively, in a consistent manner.

In order to examine how snoRBP interaction profiles compare across cell types, we performed FBL and DKC1 eCLIP in 293T and HepG2 cells in addition to K562 cells. The enrichment of snoRNAs across the profiled cell lines is highly correlated (Fig. S1E). When considering well expressed snoRNAs present at more than 10 reads per million in at least one cell type, only 6 snoRNAs (SNORD50, SNORD64, SNORD109, SNORD115, SNORD116, and SNORA35) show cell type-specific binding to FBL or DKC1 (Fig. S1F). These data suggest that the snoRNA landscape is highly consistent across cancer cell types. In summary, we comprehensively profiled the RNA interactions of all core proteins in both the C/D and H/ACA snoRNP complexes. We demonstrated that the core proteins within each complex coordinately associate with the respective C/D or H/ACA snoRNAs and that the binding specificity of snoRNP complexes is strongly cell type independent.

### Chimeric eCLIP of core C/D snoRNP proteins comprehensively recovers known C/D snoRNA interactions

Chimeric CLIP approaches enable unambiguous identification of RNA-RNA interactions by taking advantage of an intramolecular ligation of interacting RNAs during immunoprecipitation, which can then be read out as a ‘chimeric’ sequencing read that contains both fragments [15]. Prior analysis of CLIP for C/D snoRBPs successfully recovered C/D snoRNA-target interactions through mapping chimeric reads [14,15,28]. Here, we set out to determine whether we could deeply interrogate snoRNA-target interactions across cell lines and organisms with chimeric eCLIP, using core C/D snoRNP proteins as bait (Fig. 2A). Chimeric eCLIP combines the library preparation optimizations in eCLIP with an additional ligation step to encourage intermolecular ligations between co-precipitated RNA-target fragments [32]. SnoRNA targets have been extensively queried in human cancer cell lines but less characterized in rare cell types or other mammalian systems, so we profiled mouse embryonic stems cells (mESC) in addition to a standard human cell line (293T) and implemented a computational pipeline to identify snoRNA:target chimeric reads (Fig. S2A).

**Figure 2.**
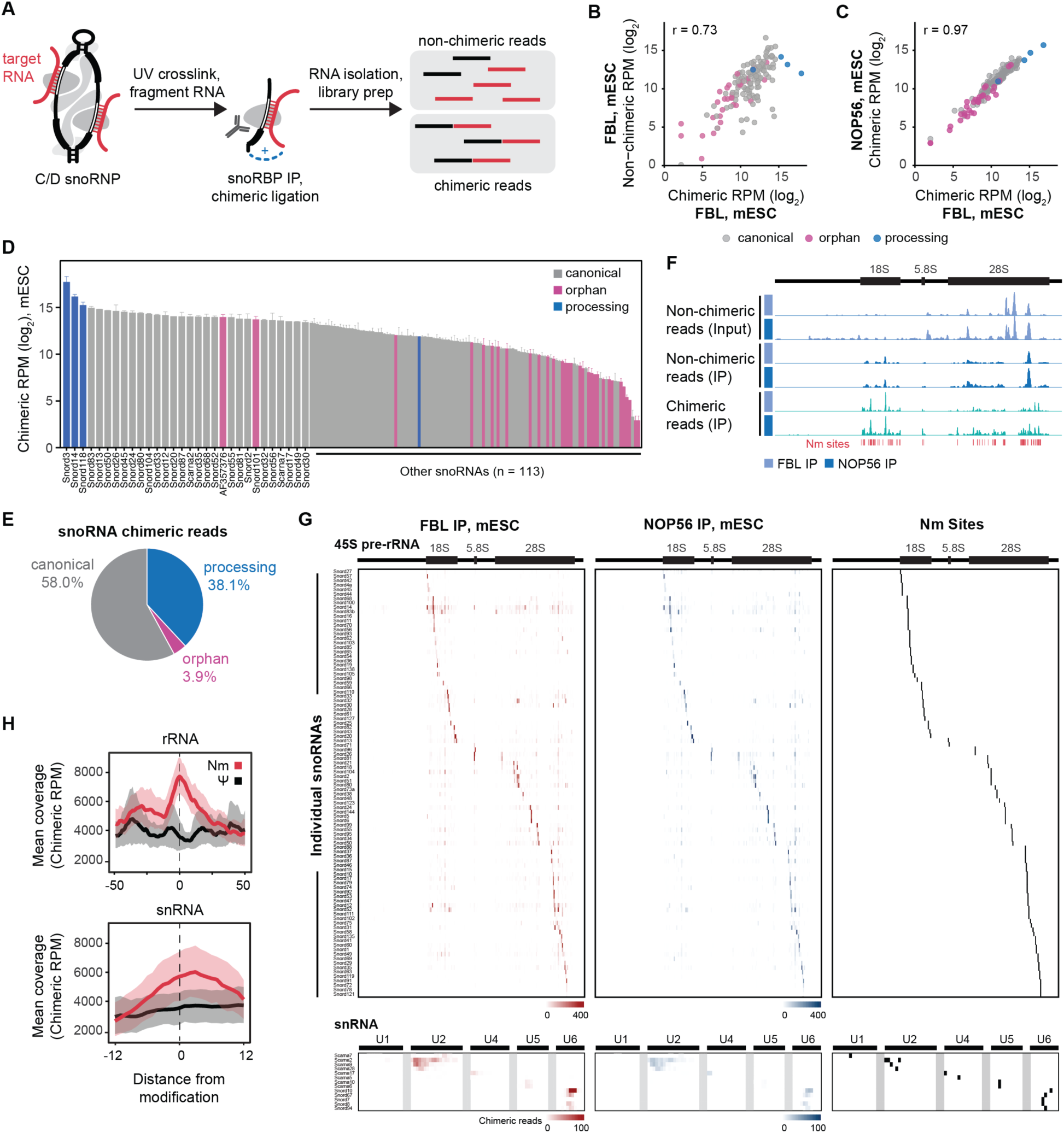
Chimeric eCLIP of core C/D snoRNP proteins comprehensively recovers known C/D snoRNA interactions in mouse embryonic stem cells (mESC). **(A)** Schematic of chimeric eCLIP procedure. **(B)** Correlation of non-chimeric versus chimeric snoRNA read abundance from FBL chimeric eCLIP. Pearson correlation (R) is indicated. **(C)** Correlation of snoRNA chimeric read abundance from FBL versus NOP56 chimeric eCLIP. Pearson correlation (R) is indicated. **(D)** Chimeric read abundance for individual snoRNAs colored according to snoRNA functional class. The y-axis indicates average of chimeric RPM in FBL and NOP56 chimeric eCLIP. **(E)** Pie chart of total number of snoRNA chimeric reads according to snoRNA functional class from FBL chimeric eCLIP. **(F)** Browser tracks of non-chimeric reads and snoRNA chimeric reads mapped to pre-rRNA as captured by FBL and NOP56 chimeric eCLIP. Annotated Nm sites are shown. **(G)** Heatmap of chimeric read coverage across pre-rRNA and snRNA from FBL and NOP56 chimeric eCLIP for canonical snoRNAs. Known Nm target sites are shown in black. **(H)** Metagene plots of chimeric read coverage flanking Nm (red) and pseudouridine (black) sites in rRNA and snRNA. RPM, reads per million.

Chimeric eCLIP generated 0.2 – 0.6% chimeric reads out of total reads in FBL and NOP56 libraries from mESC and FBL, NOP56, and NOP58 libraries from 293T cells (Fig. S2B). The efficiency of chimeric read formation was on par with or slightly better than a previously reported ligation-based approach [11]. The per-snoRNA abundance was highly correlated between non-chimeric and chimeric reads (Fig. 2B), indicating that chimeric ligation does not cause biased recovery of snoRNAs. The number of chimeric reads per snoRNA was highly correlated between different core C/D snoRBP baits in both mESC (Fig. 2C) and 293T cells (Fig. S2C). This result, similar to our observations by eCLIP, suggests a consistent recovery of snoRNA-target interactions using any of the core C/D snoRBPs. It is noteworthy that the number of chimeric reads between core C/D snoRBPs was highly correlated for SNORD13, an acetylation-guiding C/D snoRNA, as well as orphan C/D snoRNAs that lack known targets (Fig. 2C, S2C). These results implicate FBL as a constitutive component of the C/D snoRNP complex irrespective of Nm-guiding activities, and they further justify the suitability of FBL chimeric eCLIP approaches for identifying targets of orphan snoRNAs.

We captured chimeric reads for nearly all expressed C/D snoRNAs in each cell line, namely 143 out of 145 (99%) expressed snoRNAs in mESC [RPM >= 3] (Fig. 2D) and 116 out of 125 (92%) expressed snoRNAs in 293T cells [RPM >= 3] (Fig. S2D). The capture of chimeric reads to a larger proportion of snoRNAs in mESC compared to 293T cells was likely due to the greater sequence depth of the mESC libraries (Fig. S2B; 80M raw reads in mESC versus 30M raw reads in 293T). The highest numbers of chimeric reads were captured for highly abundant snoRNAs that mediate pre-rRNA processing, such as Snord3, Snord14, and Snord118, but we observed substantial coverage for both ‘canonical’ snoRNAs, which guide modification on rRNA and snRNA, and orphan snoRNAs whose targets are unknown (Fig. 2D,E, S2D).

As an initial evaluation of the success of chimeric eCLIP in identifying known C/D snoRNA-target interactions, we mapped chimeric reads to pre-rRNA and snRNA. Chimeric reads were observed primarily to the mature regions of rRNA and had a distinct distribution profile compared to non-chimeric reads from input or IP samples (Fig. 2F). Visual examination of individual snoRNA:rRNA and snoRNA:snRNA chimeric read tracks showed a marked enrichment of chimeric reads at their known Nm-guiding target sites in mESC (Fig. 2G) and 293T cells (Fig. S2E). Furthermore, metagene analysis highlighted a significant enrichment of chimeric reads at Nm sites in both rRNA and snRNA that is not observed at pseudouridylation sites (Fig. 2H, S2F). All together, these analyses lend strong support to the power and accuracy of chimeric eCLIP in experimentally detecting snoRNA-target interactions by C/D snoRBPs.

### Transcriptomic mapping of C/D snoRNA chimeras calls novel, high-confidence interactions

Next, we performed transcriptome-wide mapping of the chimeric reads obtained from core C/D snoRBP chimeric eCLIP in mESC and 293T cells. We examined the chimeric read distribution amongst different RNA classes. The majority of snoRNA chimeras mapped to rRNA (∼75%, mESC; ∼55%, 293T), and to a lesser extent, pre-mRNA (∼10%, mESC; ∼20%, 293T), snRNA (∼0.3%, mESC and 293T), and tRNA (∼0.1%, mESC; ∼0.4%, 293T) (Fig. 3A, S3A). When the chimeric read distribution was analyzed for individual canonical snoRNAs, we observed that rRNA-targeting snoRNAs had the highest proportion of rRNA-mapping chimeras, whereas some snRNA-targeting snoRNAs had an appreciably higher proportion of snRNA-mapping chimeras (Fig. S3B). Interestingly, the chimeric read distribution for individual orphan snoRNAs hinted at possible target classes; Snord23 and Snord89 chimeras were enriched for snRNAs while Snord90 and Snord101 were enriched for rRNA (Fig. S3B).

**Figure 3.**
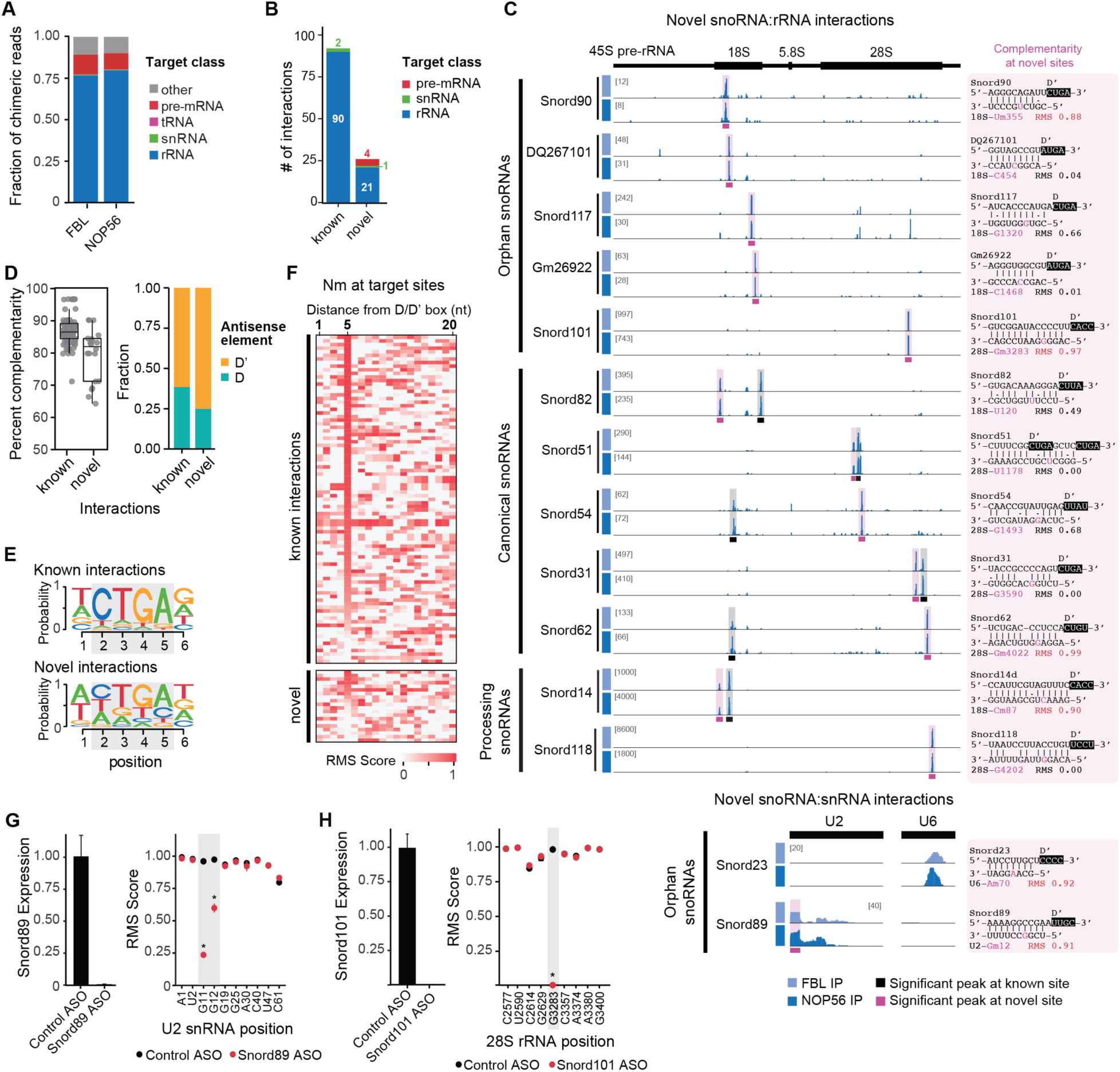
Transcriptomic mapping of C/D snoRNA chimeras calls novel, high-confidence interactions in mESCs. **(A)** Stacked bars indicate the fraction of C/D snoRNA chimeric reads mapping to distinct classes of possible target RNAs from FBL and NOP56 chimeric eCLIP. **(B)** Number of high-confidence snoRNA-target interactions identified by chimeric eCLIP according to interaction status and target class. **(C)** Browser tracks of snoRNA chimeric reads mapped to pre-rRNA or snRNA as captured by FBL and NOP56 chimeric eCLIP from mESC. Significant peaks at (black) known or (pink) novel sites. (Right) Base pair complementarity of novel interactions. RiboMeth-seq (RMS) score is shown for the 5^th^ nucleotide from the D/D’ box when high methylation is observed. **(D)** (Left) Percent complementarity of base pairing in snoRNA-target interactions for significant peaks at known or novel sites and (right) fraction of interactions associated with D or D’ antisense elements. **(E)** Consensus sequence of the D or D’ box predicted to guide the known or novel snoRNA-target interactions. **(F)** RMS score at rRNA or snRNA target sites for known versus novel interactions. The 5^th^ nucleotide is the expected position for methylation. **(G)** (Left) Snord89 expression following treatment with non-targeting (control) or Snord89-targeting ASO and (right) RMS scores at methylated nucleotides in U2 snRNA. * indicates P<0.05, two-tailed unpaired Welch’s t-test. **(H)** (Left) Snord101 expression following treatment with non-targeting (control) or Snord101-tageting ASO and (right) RMS scores at methylated nucleotides in 28S rRNA. * indicates P<0.05, two-tailed unpaired Welch’s t-test.

To call high-confidence interaction sites across the transcriptome, we performed peak calling using Clipper [33]. Peaks at known interaction sites were readily distinguished from poorly supported ‘background’ peaks when two parameters were considered: peak fold enrichment (chimeric reads from snoRBP IP relative to non-chimeric reads from input) and fraction of per-snoRNA reads in peak (chimeric reads in peak relative to total number of per-snoRNA chimeric reads) (Fig. S3C). The fraction of per-snoRNA reads in peak (i.e. the most predominant peak(s) for each snoRNA) was deemed a good classifier of positive interactions as the receiver operating characteristic curve (ROC) for known snoRNA-rRNA interactions presented an area under the curve of 99.2% in mESC and 97.4% in 293T cells (Fig. S3D).

Based on the optimal threshold determined by ROC (0.026, mESC; 0.075, 293T), we selected a peak fraction ≥ 0.1 and peak fold enrichment ≥ 3 as thresholds to minimize the false positive rate and stringently call high-confidence interactions. We called a maximum of two significant peaks per snoRNA (Fig. S3E), consistent with the presence of two antisense elements per snoRNA. When considering significant peaks called in both FBL and NOP56 chimeric eCLIP datasets from mESCs (i.e. reproducible peaks), we identified high-confidence interactions for 70% (102 out of 145) of expressed snoRNAs. This included significant peaks at 84% (90 out of 107) of known snoRNA-rRNA interactions and two known snoRNA-snRNA interactions (Fig. 3B). Peaks were present at an additional 17 known rRNA and snRNA interaction sites but fell below the stringent threshold (Fig. S3F). In 293T cells, we called a smaller percentage of snoRNA-rRNA interactions that are annotated in snoDB (Fig. S3F,G), likely reflecting the increased annotation of rare snoRNAs in human cancer cells.

Importantly, our analyses detected significant, reproducible peaks at a number of novel snoRNA interaction sites (Table S1). This included 21 novel snoRNA-rRNA and 1 novel snoRNA-snRNA interactions in mESC (Fig. 3B). While an additional 4 novel interactions were observed with pre-mRNA, the peaks were located around or adjacent to intron-embedded snoRNAs (Fig. S3H), potentially indicating spurious chimeric RNA formation during snoRNA biogenesis or incomplete snoRNA annotation. In total, we assigned high-confidence interactions for 10 orphan snoRNAs (Table S1); for example, Snord89 with U2 snRNA and Snord90, Snord101, Snord117, DQ267101, and Gm26922 with rRNA (Fig. 3C). We also observed an interaction between orphan Snord23 and U6 snRNA, but it fell below the significance cutoff (Fig. 3C). Beyond orphan snoRNAs, we called novel interactions for canonical snoRNAs that already have one antisense element assigned as an Nm guide. For example, Snord31, Snord51, Snord54, Snord62, and Snord82 each had two significant peaks in rRNA (Fig. 3C) — one peak at the known Nm target site and a second peak at the novel site. Lastly, we detected significant, reproducible peaks for two snoRNAs that mediate pre-rRNA processing, Snord14 and Snord118 (Fig. 3C). A high degree of concordance was observed across reproducible peaks in mESC and 293T cells (Table S1).

Box C/D snoRNAs base pair with their targets through an antisense element located immediately upstream of the D or D’ box element. To further characterize the interactions identified by chimeric eCLIP, we predicted the snoRNA antisense element associated with each interaction, then quantified the base pair complementarity of the snoRNA-target interaction. The base pair complementarity for novel snoRNA-rRNA interaction sites was slightly lower than known sites (78.1% mean identity at novel sites versus 86.8% at known sites) and was more commonly associated with D’ box elements (Fig. 3D). The strength of the conserved ‘CTGA’ box motif was higher for known snoRNA-rRNA interactions compared to novel interactions (Fig. 3E), which is consistent with the lower conservation of D’ box motifs compared to D box motifs. This observation suggests that the novel interactions identified by chimeric eCLIP more frequently involve less conserved D’ box motifs, which could explain why these interactions have not been identified by previous bioinformatic approaches utilizing sequence complementarity modeling.

Canonical snoRNAs guide Nm to the site on target RNA that is base paired exactly 5 nucleotides upstream of the snoRNA D/D’ box. To determine potential snoRNA-guided methylation at novel interactions in mESC, we quantified Nm levels in rRNA and snRNA by RiboMeth-seq (RMS) [34]. High levels of methylation were observed at the fifth nucleotide position for nearly all known snoRNA interactions detected in our study (mean RMS = 0.78, Fig. 3F). Novel interactions displayed overall lower RMS scores at the predicted methylation sites (mean RMS = 0.34, Fig. 3F), indicating possible methylation-independent functions of these interactions. Several novel sites, though, did display high levels of methylation at the expected nucleotide, including 18S-C87 (Snord14), 18S-U355 (Snord90), 28S-G3283 (Snord101), 28S-G4022 (Snord62), U2-G12

(Snord89), and U6-A70 (Snord23) (Fig. 3C). Snord60 is already assigned as the guide for modification at 28S-G4022, so the co-detection of Snord60 and Snord62 at this site in our study could indicate compensatory or competitive interactions that merit further investigation.

Based on our data we hypothesized that Snord89 guides methylation at U2-G12, as similarly observed in a yeast cell system [35]. To examine the targeting specificity of Snord89 in mouse cells, we performed loss-of-function experiments followed by RiboMeth-seq. Surprisingly, we found that ASO-mediated knockdown of Snord89 resulted in loss of methylation at two neighboring nucleotides in U2, as methylation at U2-G11 and U2-G12 were decreased by 72% and 37%, respectively (Fig. 3G). We then validated the targeting specificity of Snord101 using the same approach. ASO-mediated knockdown of Snord101 resulted in complete loss of methylation at 28S-G3283 with no effect on the flanking Nm sites (Fig. 3H).

We also tested whether the chimeric eCLIP approach could capture H/ACA snoRNA-target interactions by performing chimeric eCLIP with DKC1 and GAR1 as the bait proteins from 293T cells. Analysis of non-chimeric reads from chimeric eCLIP yielded similar results to standard eCLIP, as over 24% of reads mapped to snoRNAs with greater recovery for H/ACA snoRNAs ( 16% and 20% of total reads by DKC1 and GAR1) versus C/D snoRNAs (7% in both RBPs) (Fig. S4A). Chimeric analysis recovered ∼1% chimeric reads out of total reads, but despite the higher recovery of H/ACA snoRNAs among non-chimeric reads, we observed that less than half of the chimeric reads contained an H/ACA snoRNA (Fig. S4B). Manual inspection of well-studied H/ACA snoRNAs suggested that while some known sites may be captured, most showed poor recovery and resolution at known target regions (Fig. S4C). Performing peak calling confirmed that DKC1 chimeric eCLIP had at best marginal recovery of true interactions (Fig. S4D). We hypothesize that the distinct structural complexity of H/ACA snoRNAs may make them refractory to standard chimeric CLIP approaches and that alternative approaches will be required to profile H/ACA interactomes.

Thus, these results confirm that chimeric eCLIP can be used to find new C/D snoRNA interactions for but requires further optimization for H/ACA snoRNAs. We identified high-confidence interactions with reproducibility between different core C/D snoRBP baits in rRNA and snRNA, a subset of which may involve alternative functions or regulatory mechanisms beyond 2’-O-methylation. Compared to bioinformatic approaches, this experimental approach is agnostic to the identification of D/D’ box motifs, target RNA class, or presence of chemical modification at the target site.

### Identification of specific subsets of snoRNA-target interactions using orthogonal protein baits

In addition to the core C/D and H/ACA snoRNA binding proteins, several additional RBPs have been reported to associate with snoRNAs and regulate their functions beyond guiding rRNA modifications [24,36]. As these interactions are under-explored, we set out to test whether performing chimeric eCLIP with less-studied snoRNA-associated proteins would enrich for functionally related subsets of snoRNAs and help uncover new aspects of snoRNA biology. We first identified candidate bait proteins to use in our study by integrating our core C/D and H/ACA snoRBP eCLIP datasets with the 150 RBPs profiled by ENCODE to analyze snoRNA enrichment with a customized analysis pipeline [29,37]. While the majority of evaluated RBPs displayed a depletion for both C/D and H/ACA snoRNAs, 11 RBPs (in addition to the core snoRBPs above) showed at least 4-fold enrichment for snoRNAs and may represent potential snoRNA partners (Fig. 4A). These non-core snoRBPs clustered into two functional annotations: a cluster of RBPs that included splicing- and spliceosomal RNA-associated factors (LARP7, NOLC1, PTBP1, SMNDC1, TRA2A, and ZC3H8), discussed further below, and a cluster of rRNA processing factors (AATF, UTP3, UTP18, WDR3, WDR43) which showed even stronger specificity for C/D snoRNAs than the core C/D snoRBPs (Fig. 4A). Considering individual snoRNA enrichment, we observed that these rRNA processing RBPs, members of the small-subunit (SSU) processome [38,39], were particularly enriched for the processing snoRNA SNORD3 (U3) relative to FBL (Fig. 4B, S5A). Comparing SNORD3 fold-enrichment across all RBPs in ENCODE confirmed that rRNA processing RBPs, and particularly the small-subunit processome, showed strong enrichment for SNORD3 (Fig. 4C, S5B), consistent with the critical role of SNORD3 in coordinating the function of the SSU in rRNA processing [40].

**Figure 4.**
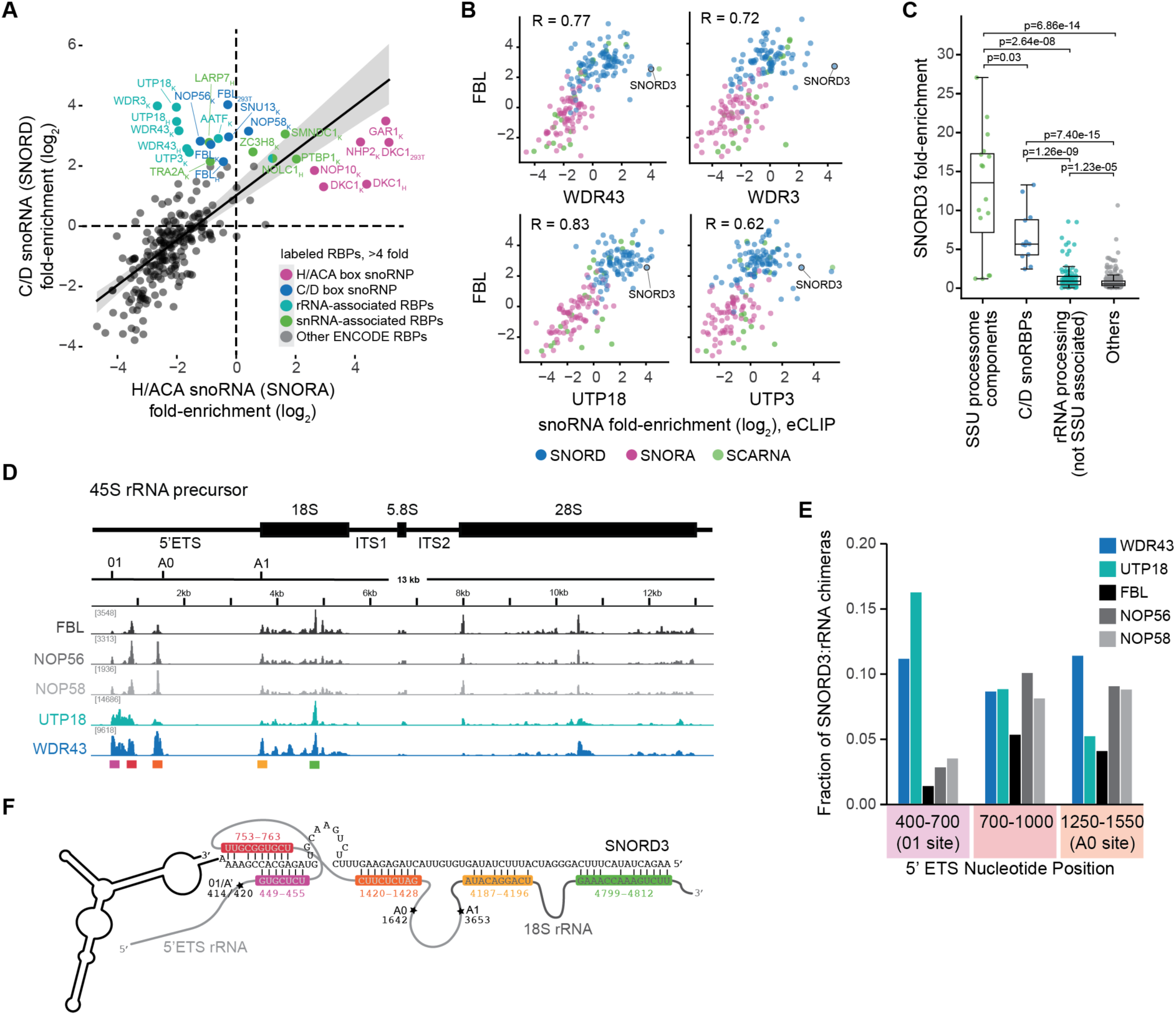
RBPs that function in ribosomal RNA processing recover unique SNORD3 chimeric reads. **(A)** Fold-enrichment correlation between (x-axis) H/ACA and (y-axis) C/D snoRNAs across 8 snoRBPs and 223 ENCODE eCLIP datasets. **(B)** Correlation of snoRNA fold-enrichment in FBL and ribosomal RNA-associated RBPs in K562 cells. correlation (R) is shown. Only snoRNAs with 5 or more reads in IP were plotted. **(C)** Box plot showing fold-enrichment of SNORD3 in eCLIP datasets for indicated groups of RBPs. Two-sample Kolmogorov-Smirnov (KS) test was performed. **(D)** Browser tracks of SNORD3 chimeric reads mapped to pre-rRNA as captured by chimeric eCLIP for the indicated RBPs. **(E)** Bars indicate the fraction of SNORD3 chimeric reads at different processing sites in the 5’ETS region of pre-rRNA (relative to total SNORD3:rRNA chimeric reads). **(F)** Schematic of predicted structures of potential SNORD3-rRNA interactions, with labelled regions colored according to (D).

To test whether we could recover distinct interaction profiles of SNORD3 associated with these SSU factors versus the core C/D snoRNP complex, we performed chimeric eCLIP for WDR43 and UTP18 in K562 cells (Fig. S5C). Although SNORD3 was typically the most frequently observed snoRNA in chimeric reads for all RBPs profiled (Fig. S5D), WDR43 and UTP18 showed an even higher frequency of SNORD3:rRNA chimeras compared to FBL, NOP56, or NOP58 (Fig. S5E). Visual examination of SNORD3:rRNA chimeric read tracks showed a strong enrichment of SNORD3 chimeric reads in the 5’ETS region of pre-rRNA, with a distinct interaction profile for UTP18 and WDR43 compared to FBL or other core C/D snoRBPs (Fig. 4D, purple, red, and orange peaks). SNORD3 chimeric reads were enriched and preferentially positioned at the 01-A0 early processing sites with WDR43 and UTP18 pulldown compared to FBL (Fig. 4E, S5F), consistent with SSU-associated SNORD3 playing a key role in interacting with these sites during early ribosomal processing. We also see capture of SNORD3 chimeras with UTP18 and WDR43 at other downstream sites in pre-rRNA in a manner distinct from FBL, suggesting that SSU-mediated SNORD3 interactions may also drive later rRNA processing steps (Fig. 4D, yellow and green peaks). Notably, sequence alignment indicates that sequences within the 01/A’ (400-700 nucleotide) and intermediate 700-1000 nucleotide regions of pre-rRNA are complementary to overlapping regions of SNORD3 (Fig. 4F), suggesting that the chimeras may reflect sequential interactions during the complex series of processing steps involved in pre-rRNA transcription, folding, and maturation [41]. Overall, these results indicate that chimeric eCLIP for SSU processome components can recover SNORD3 interactions distinct from those observed with FBL, validating the principle that using unique protein baits can reveal new snoRNA interaction landscapes.

### Chimeric eCLIP of LARP7 and NOLC1 detects unique snoRNA-snRNA interactions

Post-transcriptional modifications of snRNAs are essential for spliceosomal fidelity and are largely guided by snoRNAs, with the scaRNA class playing a particularly important role [42,43]. More snoRNA-snRNA interactions continue to be discovered, including our identification of Snord23-U6 and Snord89-U2 interactions above and other recent work [18]. This, together with a recent study that identified LARP7 as a novel bridging factor for snoRNA-guided modifications on U6 [25,44], suggests that more principles of snRNA regulation by snoRNAs are yet to be uncovered. As such, we next asked whether chimeric profiling of the splicing- and spliceosomal RNA-associated proteins we identified above could enrich for unique snoRNA-snRNA regulatory networks.

As validation of this approach, we first examined LARP7. Previous studies proposed a mechanism in which LARP7 binds the subset of C/D snoRNAs that have been shown to guide U6 2’-O-methylation, including SNORD7, SNORD8, SNORD9, SNORD10, SNORD67, and SNORD94 [25]. Consequently, we performed chimeric eCLIP on LARP7 and FBL in HepG2 cells (Fig. S6A) to determine if LARP7 could capture and enrich for these snoRNA-U6 interactions. We observed that the previously characterized LARP7-associated snoRNAs had the highest enrichment among non-chimeric (i.e. snoRNA only) and chimeric reads in LARP7 compared to FBL chimeric eCLIP (Fig. 5A,B, S6B). SnoRNA:U6 chimeras were recovered for five of the six LARP7-associated snoRNAs, and these chimeras mapped to the known U6 Nm target sites in both FBL and LARP7 datasets (Fig. 5C,D), validating the role of LARP7 in bridging functional snoRNA-snRNA interactions. Many of the other U6-interacting snoRNAs annotated in snoDB were not captured by FBL, and manual inspection of their annotation source found these were often supported by sparse interaction data and lacked validation of snoRNA-responsive methylation. Overall, recovery of chimeras for known snoRNA-U6 interactions was 2.2 times higher in LARP7 versus FBL chimeric eCLIP, whereas recovery of chimeras for known snoRNA-rRNA interactions was dramatically lower (11.9 times) (Fig. 5E,F). These data confirm that chimeric eCLIP can capture and enrich for specific subsets of snoRNA interactions involving LARP7.

**Figure 5.**
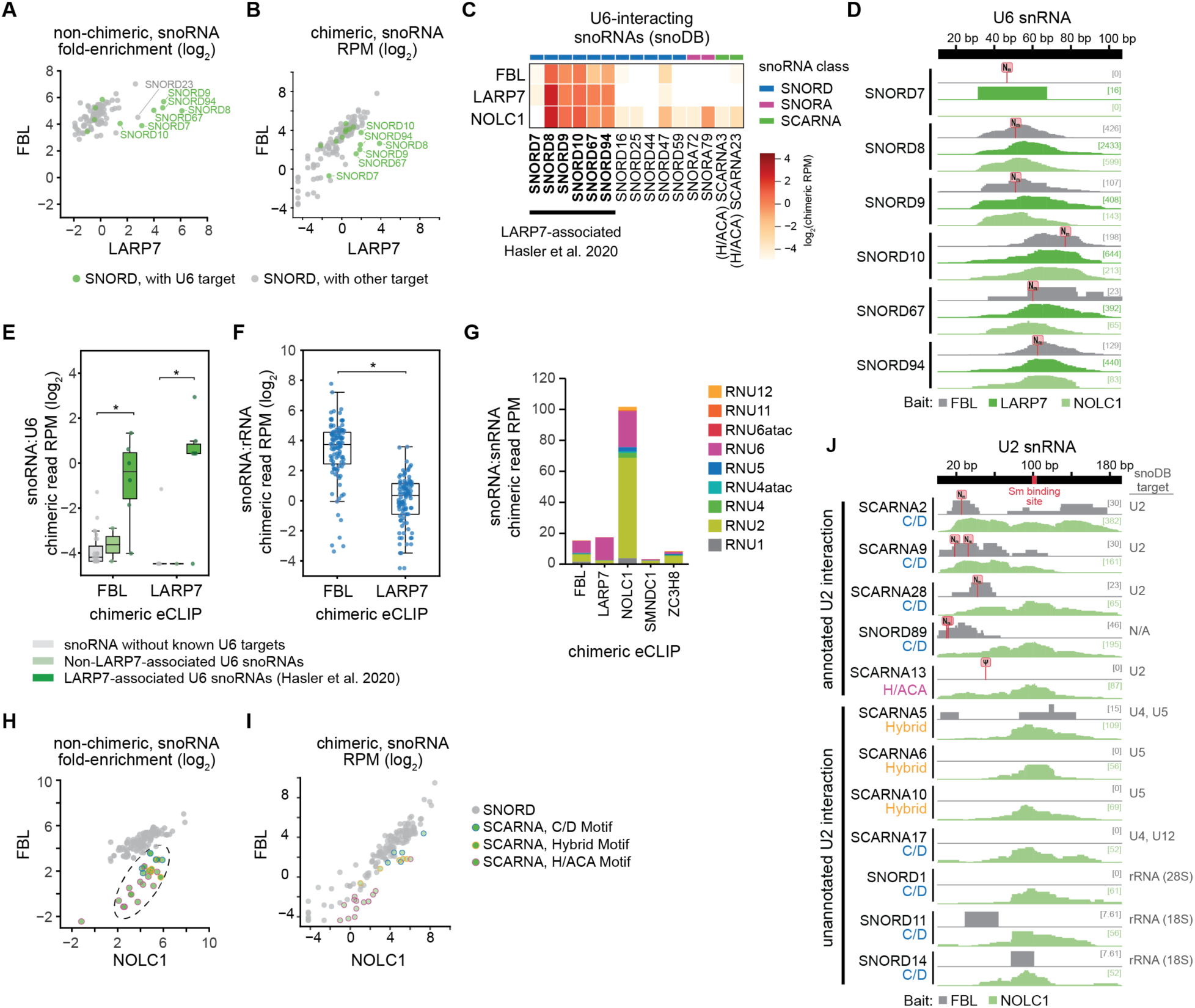
Chimeric eCLIP for spliceosomal RNA-associated RBPs enrich for snoRNA-snRNA interactions. **(A)** Points indicate C/D box snoRNA fold-enrichment of non-chimeric reads from chimeric eCLIP of FBL versus LARP7 in HepG2 cells. Datapoints for C/D snoRNAs with U6 interactions annotated in snoDB are colored. Datapoints with less than 5 reads in IP were discarded. **(B)** Points indicate C/D box snoRNA chimeric read abundance (RPM) from chimeric eCLIP of FBL versus LARP7. Datapoints for C/D snoRNAs with U6 interactions annotated in snoDB are colored. **(C)** Heatmap of snoRNA chimeric read abundance (RPM) from FBL, LARP7, and NOLC1 chimeric eCLIP for all snoRNAs with U6 interactions annotated in snoDB. **(D)** Browser tracks show read density (per million chimeric reads) for snoRNA chimeras along the U6 snRNA for U6-interacting snoRNAs. **(E)** Points indicate snoRNA:U6 chimeric read abundance (RPM) from FBL or LARP7 chimeric eCLIP. Colors indicate snoRNAs classified as (green) LARP7-associated U6 snoRNAs, (light green) Non-LARP7-associated U6 snoRNAs, and (grey) all other snoRNAs. * indicates P < 0.05, unpaired two sample t-test. **(F)** Points indicate snoRNA:rRNA chimeric read abundance (RPM) for snoRNAs with known rRNA target sites from FBL or LAPR7 chimeric eCLIP. * indicates P < 0.05, unpaired two-sample t-test. **(G)** Stacked bar plot indicates the target frequency of snoRNA:snRNA chimeric reads (RPM) recovered in chimeric eCLIP for the indicated RBPs. **(H)** Points indicate non-chimeric snoRNA fold-enrichment in FBL versus NOLC1 chimeric eCLIP in HepG2 cells. Datapoints with less than 5 reads in IP were discarded. Circle outline color represents the type of SCARNA with (blue) C/D box motif, including C/D SCARNAs and Tandem-CD SCARNAs), (pink) H/ACA box motif, including H/ACA SCARNAs and Tandem-H/ACA SCARNAs, and (orange) hybrid SCARNAs that possess both C/D and H/ACA box motifs. **(I)** Points indicate snoRNA chimeric read abundance (RPM) in FBL versus NOLC1 chimeric eCLIP. **(J)** Browser tracks show read density (per million chimeric reads) for snoRNA:U2 chimeric reads along the U2 snRNA for known and candidate U2-interacting snoRNAs. RPM: reads per million sequenced reads.

We then looked for novel RBP baits that could highlight additional snoRNA-snRNA interaction principles. We further analyzed the ENCODE RBP eCLIP data and identified four spliceosomal RNA-associated proteins (NOLC1, SMNDC1, ZC3H8, and PTBP1) with co-enrichment for both snoRNAs and scaRNAs (Fig. S6C). These RBPs were also often enriched for either U2 or U6 snRNAs, echoing their known roles in snRNA processing (Fig. S6D,E) [45–48]. Next, we performed chimeric eCLIP for NOLC1, SMNDC1, and ZC3H8 in the same cell lines as the ENCODE eCLIP experiments: NOLC1 in HepG2 cells and SMNDC1 and ZC3H8 in K562 cells (Fig. S6A). Due to technical issues, PTBP1 was not profiled. We noted that the size-resolved band of NOLC1 was present in standard eCLIP but lost after chimeric ligation (Fig. S6A,F), suggesting that this ligation may create a mixture of higher-order complexes.

NOLC1 chimeric eCLIP enriched for snoRNA:snRNA chimeras compared to either FBL or LARP7 (Fig. S6G), with a pronounced enrichment for both U2 and U6 chimeras (Fig. 5G). In contrast, SMNDC1 and ZC3H8 had low recovery of snoRNA:snRNA chimeras for U2 and U6 despite their enrichment among non-chimeric reads (Fig. 5G, S6H), and we did not pursue further analysis on these RBPs. Unlike LARP7, NOLC1 chimeric eCLIP did not enrich for a specific subset of C/D snoRNAs; rather, the entire class of scaRNAs was enriched in NOLC1 versus FBL chimeric eCLIP among both non-chimeric reads (Fig. 5H) as well as chimeric reads (Fig. 5I). This included not only scaRNAs containing box C/D motifs, but also H/ACA and hybrid motifs, consistent with the role of NOLC1 in maturation and function of both snoRNA classes [49].

For C/D-containing snoRNAs known to target U2 or U6, in many cases NOLC1 recovered chimeras at their known target sites, as illustrated for SCARNA2 and SCARNA9 in U2 (Fig. 5J) and SNORD8, SNORD9, and SNORD10 in U6 (Fig. 5D). The enrichment profiles for these known snoRNA guides were very similar to chimeric eCLIP of FBL (Fig. 5D,J), suggesting NOLC1 may act in concert with the core snoRNP at these target sites. In addition, we also observed chimeras with snoRNAs that have not previously been linked to U2 (n = 22) and U6 (n = 9) (Fig. S6I). Interestingly, we noted that many of candidate interactions lack chimeric support in FBL pulldown, had a stereotypical positioning of NOLC1 chimeras in the central region of U2 which overlaps the Sm binding region, and contained snoRNAs that have been shown to target modifications on other snRNAs (Fig. 5J, S6J), suggesting that these unique chimeras in NOLC1 pulldown may reflect the more general role of NOLC1 in snRNA biogenesis [49,50]. In summary, these results suggest that chimeric eCLIP can successfully enrich for previously characterized snoRNA interactions, as well as provide further depth to identify and understand novel interactions driven by non-canonical snoRNP complexes.

### SNORD89 mediates U2 processing to fine-tune splice site recognition

To test whether integration of NOLC1 and FBL chimeric experiments could reveal biological insights into U2 function, we performed further studies of one poorly characterized snoRNA with strong chimeric signal in NOLC1 and FBL experiments, SNORD89. While NOLC1 captured SNORD89:U2 chimeras throughout U2 (with an enrichment in the central Sm binding region), FBL and NOP56 captured SNORD89:U2 chimeras primarily at the 5’ end of U2 (Fig. 3C, 5J), overlapping the G11 and G12 Nm target sites that are responsive to SNORD89 knockdown (Fig. 3G). The functional impact of U2 modification at these specific nucleotides remains unclear: methylation at G12 was shown to be essential for splicing progression on a single pre-mRNA *in vitro* [43] but has not been explored on transcriptome-wide *in vivo*, and the role of G11 methylation is unknown. Thus, we next wanted to examine the functional impact of SNORD89 loss on spliceosomal activity.

To quantify the effect of SNORD89 on splicing in human cells, we performed SNORD89 ASO knockdown in 293T cells (Fig. 6A), followed by RNA-seq. Analysis of alternative splicing identified 300 cassette exons more excluded upon SNORD89 knockdown (‘knockdown-excluded exons’) and 334 cassette exons more included upon SNORD89 knockdown (‘knockdown-included exons’) (Fig. 6B), confirming that SNORD89 significantly alters splicing. In further analysis, we noted that SNORD89 preferentially regulates exons with nearly constitutive splicing — 60% of knockdown-excluded exons had percent spliced in (PSI) values of 0.95 or higher in control samples, and 66% of knockdown-included exons had PSI values of 0.95 or higher upon SNORD89 ASO (Fig. 6C).

**Figure 6.**
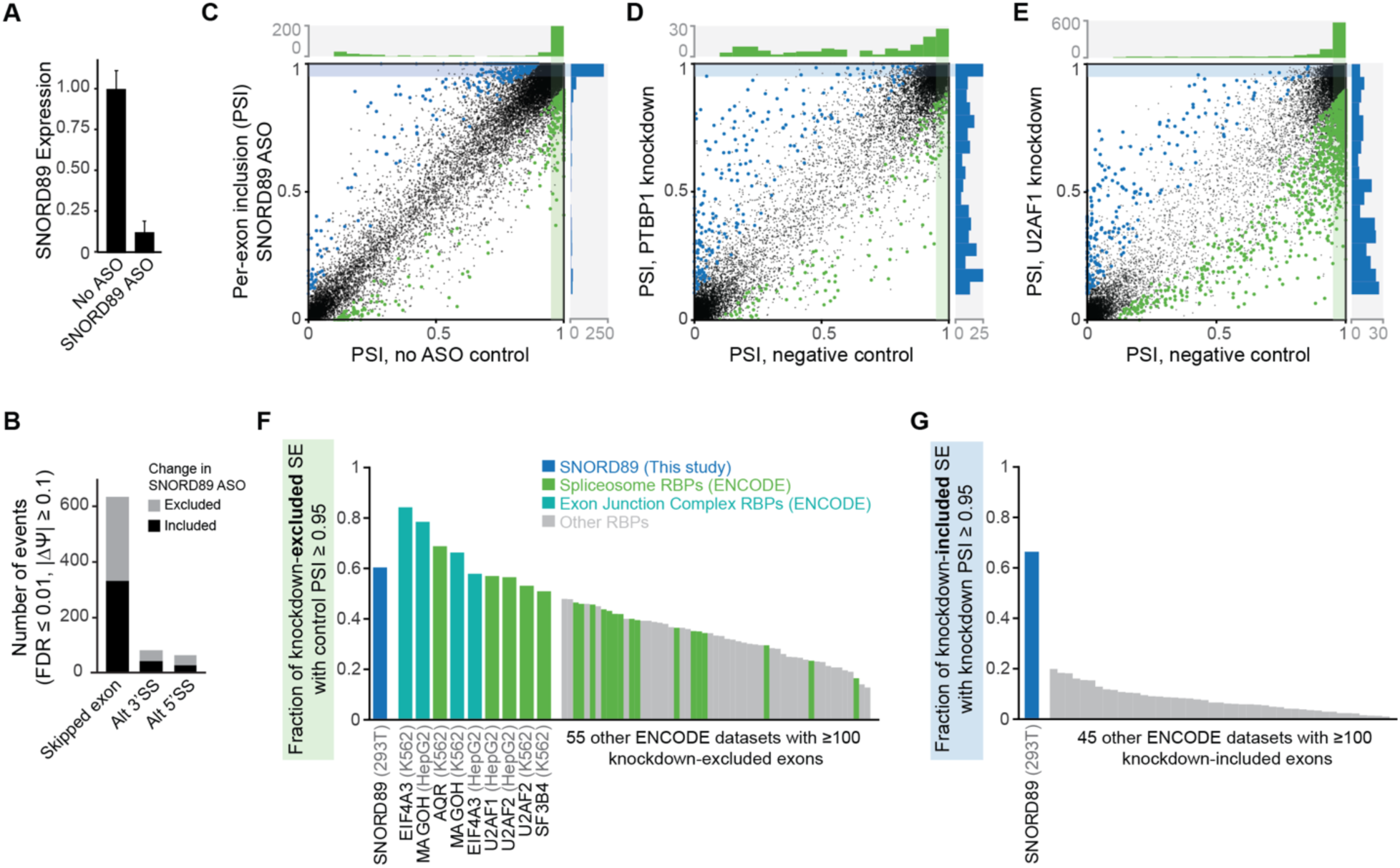
SNORD89, a guide for U2 2’-O-methylation, plays a unique role in splicing control. **(A)** Bars indicate SNORD89 expression (from RT-qPCR) upon SNORD89 antisense oligonucleotide knockdown. **(B)** Bars indicate the number of significantly altered alternative splicing events from rMATS (FDR ≤ 0.01, change in percent spliced in (|ΔΨ|) ≥ 0.1, with ≥ 10 junction reads in all samples), comparing SNORD89 ASO versus no ASO control. **(C-E)** Scatter plots indicate percent spliced in (Ψ) for (black) all exons, (green) knockdown-excluded, and (blue) knockdown-included cassette exons for (C) SNORD89, (D) PTBP1, and (E) U2AF1 knockdown. PTBP1 and U2AF1 data were obtained from the ENCODE project. Histograms indicate the frequency of Ψ values (in 0.05 bins) for (green) the control sample for knockdown-excluded exons, or (blue) the knockdown sample for knockdown-included samples. **(F-G)** Bars indicate the percent of events with Ψ ≥ 0.95 for (F) the control sample for knockdown-excluded exons, or (G) the knockdown sample for knockdown-included exons. Shown are (leftmost bar, blue) SNORD89 knockdown and (all other bars) all ENCODE RBP knockdown datasets with at least 100 significantly altered exons.

To consider whether the effect of SNORD89 on knockdown-excluded exons with high PSI values was unique, we performed re-analysis of 473 RBP knockdown datasets generated by the ENCODE consortium [29]. We observed that canonical alternative splicing regulators showed a more varied distribution of PSI values for knockdown-excluded exons whereas spliceosomal components showed a pattern similar to SNORD89. For example, only 20% (28 out of 143) of PTBP1-dependent exons had PSI ≥ 0.95 in controls (Fig. 6D), but 57% (571 out of 1003) of U2AF1-dependent exons had PSI ≥ 0.95 in controls (Fig. 6E). Indeed, considering all 64 ENCODE datasets with at least 100 knockdown-excluded exons, the observation that more than half of knockdown-excluded exons had PSI ≥ 0.95 in controls was only seen for RBPs that were core components of the spliceosome or exon junction complex (Fig. 6F). Thus, SNORD89 enables inclusion of a set of near-constitutive exons similar to core splicing machinery RBPs.

In contrast, similar analysis of knockdown-included exons across ENCODE datasets indicated that the preferential regulation of near-constitutive exons was a unique property of SNORD89, as no RBP knockdown from ENCODE showed more than 20% of knockdown-included exons with PSI ≥ 0.95 (Fig. 6G). Thus, rather than simply being essential for enabling recognition of constitutive splice sites, SNORD89-mediated modifications on U2 appear to play a more precise role in limiting complete processing of near-constitutive sites. Analysis of intron retention suggested a similar role: SNORD89 knockdown caused a 24.7% decrease in the overall fraction of intronic reads observed (from 7.4% in control to 5.6% in knockdown) (Fig. S7A), and intron-specific analysis of SNORD89 knockdown showed only 250 retained introns with increased retention but 800 retained introns with increased splicing (Fig. S7B).

The observation that SNORD89 knockdown more often led to more efficient splicing of retained introns, coupled with SNORD89 knockdown causing both increased and decreased mis-splicing of near-constitutive cassette exons, suggested that SNORD89 might be playing a unique role in maintaining both accurate and inaccurate splicing of specific introns. To test this hypothesis, we queried whether SNORD89 knockdown altered the accuracy of utilizing known splice junctions. We observed that SNORD89 knockdown caused a 12% decrease in the frequency of unannotated splice junctions (Fig. S7C), indicating that SNORD89 is acting to maintain rather than suppress utilization of unannotated junctions. In sum, these results suggest that modifications directed by SNORD89 are not only essential for proper splicing but may also be playing a critical role in maintaining splicing errors at a subset of splice sites.

## Discussion

SnoRNAs play crucial roles in the modification and processing of rRNA and snRNA, impacting fundamental cellular processes such as translation and splicing. Despite clear biological relevance, research efforts have failed to identify targets for many orphan snoRNAs, and the roles of snoRNA-associated proteins are not well characterized. To address these gaps in understanding, we implemented an improved chimeric eCLIP approach that enables deep and accurate profiling of snoRNA interactions in an RBP-dependent manner. This new approach resulted in increased depth of snoRNA:target chimeras relative to previous efforts to directly map snoRNA interactions [14,15,18,28]. The high accuracy achieved (AUC > 97% in both human and mouse cell lines) allows for benchmarking against known targets and the characterization of novel interactions. Notably, this accuracy was achieved empirically without needing to model interaction complementarity as in previous work [28], enabling recovery of candidate interactions that may have non-canonical interaction dynamics.

Using stringent peak calling criteria, we identified novel snoRNA interactions, including those of orphan snoRNAs. For example, we called a high-confidence interaction between Snord101 and 28S rRNA as well as Snord89 and U2 snRNA. While similar interactions were detected in previous chimeric ligation sequencing approaches, the predicted Nm sites were either inaccurate or lacked experimental validation [13,18]. We confirmed the bona fide effect of Snord101 and Snord89 on guiding Nm modification at 28S-G3283 and U2-G11/G12, respectively, in snoRNA loss-of-function experiments. Additionally, we identified putative interaction sites for Snord14, Snord23, Snord62 and Snord90, each supported by high methylation scores measured by RiboMeth-seq at the expected targeted nucleotide. Some of these interactions also have experimental support in other recent studies [13,14]. We also refine previous predictions; previous work that identified SNORD23:U6 chimeras proposed a candidate methylation site (U6-Um64) that has not yet been observed, whereas our analysis indicates D’-box positioning at U6-Am70, a previously observed LARP7-dependent methylation event [25]. We observed similar results across all core C/D snoRBPs, confirming the robustness of this approach.

Novel interactions showed increased use of less conserved D’ box-associated antisense elements. For example, the interaction identified between Snord89 and U2 snRNA is associated with the D’ box, featuring a ‘UUGC’ motif that diverges from the canonical ‘CUGA’ motif. Our observation that loss of Snord89 affects Nm levels at two neighboring nucleotides, including G11 which was previously predicted as a Scarna2 target site [51] and G12 which was shown to be a Snord89 target site in yeast [35], suggests that perhaps the poorly conserved D’ box motif may support more flexibility in the targeted nucleotide.

Modifications at the 5’ end of U2 snRNA are crucial for snRNP assembly and splicing [43,52]. However, their exact functions remain unclear, in part due to a lack of appropriate systems and tools. Further analysis indicated a unique role for SNORD89 in modulating spliceosome activity (Fig 6). SNORD89 knockdown caused increased skipping of many normally constitutive exons, consistent with the previously described essential role for U2-G12 in splicing [43]. Surprisingly, we also observed 222 splicing events that were near-constitutive only upon SNORD89 knockdown, suggesting that these methylations also act to inhibit splicing of otherwise efficiently recognized exons. This pattern was not observed in any of 433 RBP knockdowns performed by the ENCORE project, suggesting these modifications may play a relatively unique role in splicing control. How altering U2-G11 and U2-G12 modification state could both mimic essential spliceosomal proteins as well as uniquely suppress inclusion of near-constitutive exons is unclear, but their position suggests a possible mechanism through a role in spliceosomal structure. Prior to the first transesterification step of splicing, U2 and U6 form intra- and inter-molecular structures through base-pairing interactions that facilitate active site formation and prime the spliceosome for nucleophilic attack [53–55]. The U2:U6 interactions can take on two different conformations, a three-way junction (3WJ) or a four-way junction (4WJ), and U2-G11 and U2-G12 are strategically positioned at the interface where the 3WJ or 4WJ structures forms [56–59]. Although the functional significance of these structures for splicing remains controversial, SNORD89 methylation of either or both U2-G11 and U2-G12 could be a novel mechanism through which small RNAs and RNA methylation can fine-tune splicing kinetics by mediating the intermediate spliceosome steps that occur stepwise prior to catalytic activation [58–60]. Further careful biochemical work will be required to explore the dynamics of structures, and how U2-G11 and U2-G12 modification alter these kinetics to drive splicing changes.

Beyond the core snoRBPs, numerous studies have indicated that additional accessory RBPs can interface with snoRNAs to modulate their targets or functions [24,26]. Using snoRNA-associated accessory proteins as baits in chimeric eCLIP, we confirmed that WDR43 and LARP7 associate with a specific subset of snoRNA interactions. We observed U6 interactions with the specific snoRNAs through both LAPR7 and FBL, validating the previously model that LARP7 bridges U6 modifications [25,44]. We also observed SNORD3 interaction at various processing sites within the 5’ ETS regions of 45S pre-rRNA using core and non-core snoRBPs and these interactions illustrate how SNORD3 might arrange pre-rRNA folding via multiple interaction sites during early rRNA processing. Beyond that, we observed FBL-independent U2 interactions through accessory proteins, NOLC1. NOLC1 pulldown captured novel interactions at stereotypical positions (Sm binding sites) in both U2 and U6, suggesting these interactions may not drive canonical modifications. Although we do not yet know the function of these novel pairings, it is possible that they function as intermediates in enabling proper localization and targeting of these snoRNAs to their functional targets. Further experiments are required to understand the functions of such interactions on either U2 or other RNA molecules.

Although we successfully captured C/D snoRNA interactions by chimeric eCLIP, we were less successful at capturing H/ACA snoRNA interactions. Since we observed that eCLIP of DKC1 and other H/ACA snoRBPs successfully enriched for both H/ACA snoRNAs and (pre- and mature) rRNA, the inefficient capture of H/ACA snoRNA:target chimeras likely reflects a technical challenge of chimeric ligation. H/ACA snoRNAs contain longer stretches of both inter- and intra-molecular double-stranded RNA structure that may impair the flexibility necessary to support not only chimeric ligation, but also UV crosslinking and RNA fragmentation. We anticipate that the continued development of additional crosslinking reagents, RNA ligase enzymes, and reaction conditions will likely allow for successful capture of H/ACA interactions with a similar methodology. Our chimeric eCLIP approach currently lacks identification of non-rRNA/snRNA targets, which could likely be improved by implementing rRNA depletion during library preparation — a strategy successfully employed in previous studies [10,13]. Despite this limitation, our study successfully identified numerous novel interactions. These findings underscore the robustness of the chimeric eCLIP method and its ability to uncover significant snoRNA-target interactions, offering valuable insights into snoRNA functionality. Moving forward, future studies could also explore the use of different biological tissues or conditions to determine if snoRNA interactions vary across different biological contexts, providing a more comprehensive view of snoRNA functions.

## Supporting information

Supplemental Protocol 1

---

**Figure S1.**
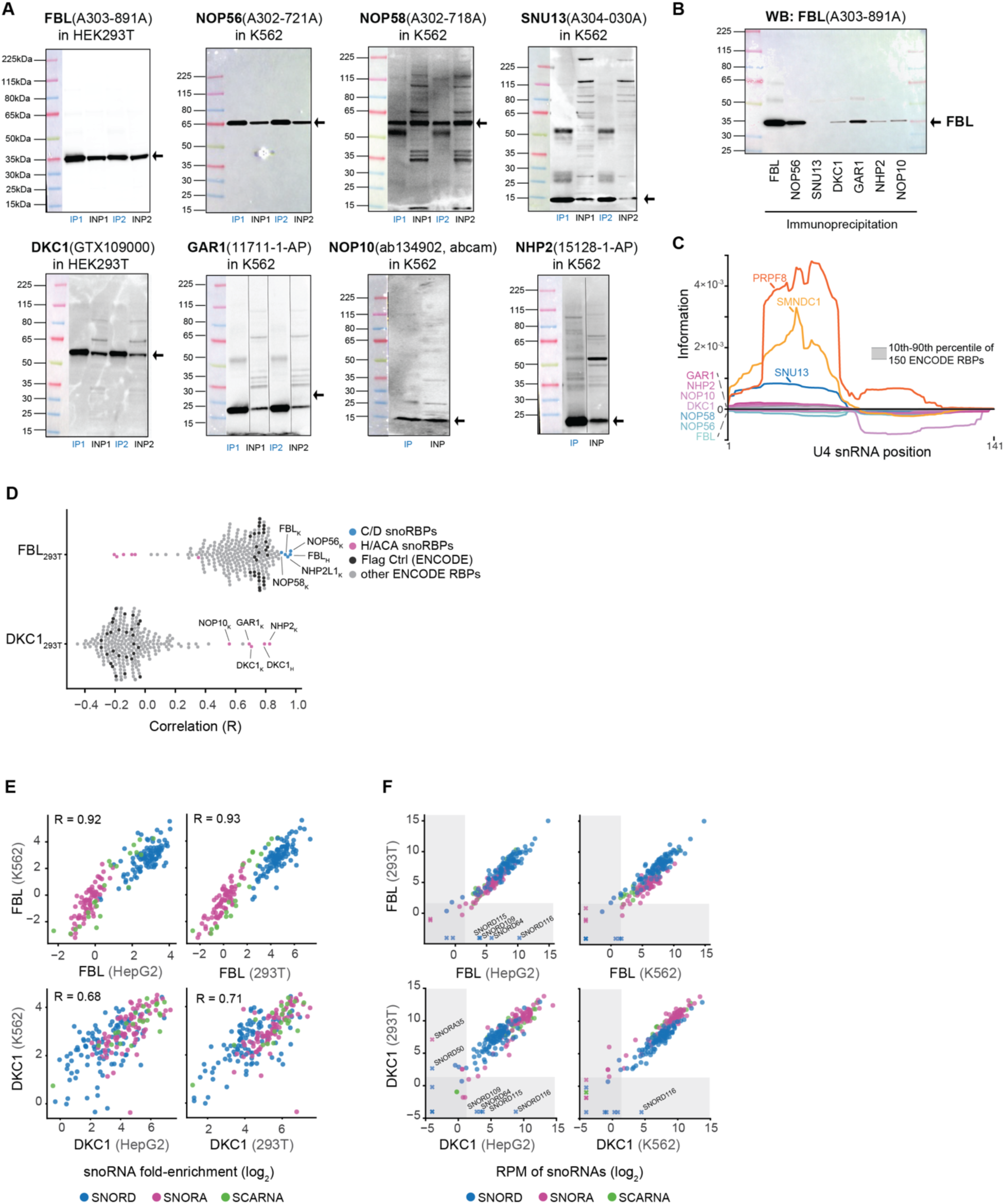
snoRNP proteins comparison in snoRNA enrichment. **(A)** Western blot confirmation of snoRBP immunoprecipitation (IP) during eCLIP in K562 cells. INP, input. **(B)** Western blot for FBL following immunoprecipitation of core snoRBPs during eCLIP. **(C)** eCLIP relative information along U4 snRNA for (colors) indicated RBPs and (grey) all ENCODE RBPs. **(D)** Pearson correlation coefficient of snoRNA fold-enrichment between all profiled RBPs and (top) FBL or (bottom) DKC1 in 293T cells. Subscripts indicate cell lines; K562 (K) and HepG2 (H). **(E)** Correlation of snoRNA fold-enrichment across (top) FBL or (bottom) DKC1 eCLIP in different cell lines. Pearson correlations (R) are indicated. **(F)** snoRNA abundance (RPM, reads per million sequenced reads) across (top) FBL or (bottom) DKC1 eCLIP in different cell lines. Data points symbolized with an ‘x’ are missing values in one cell line.

**Figure S2.**
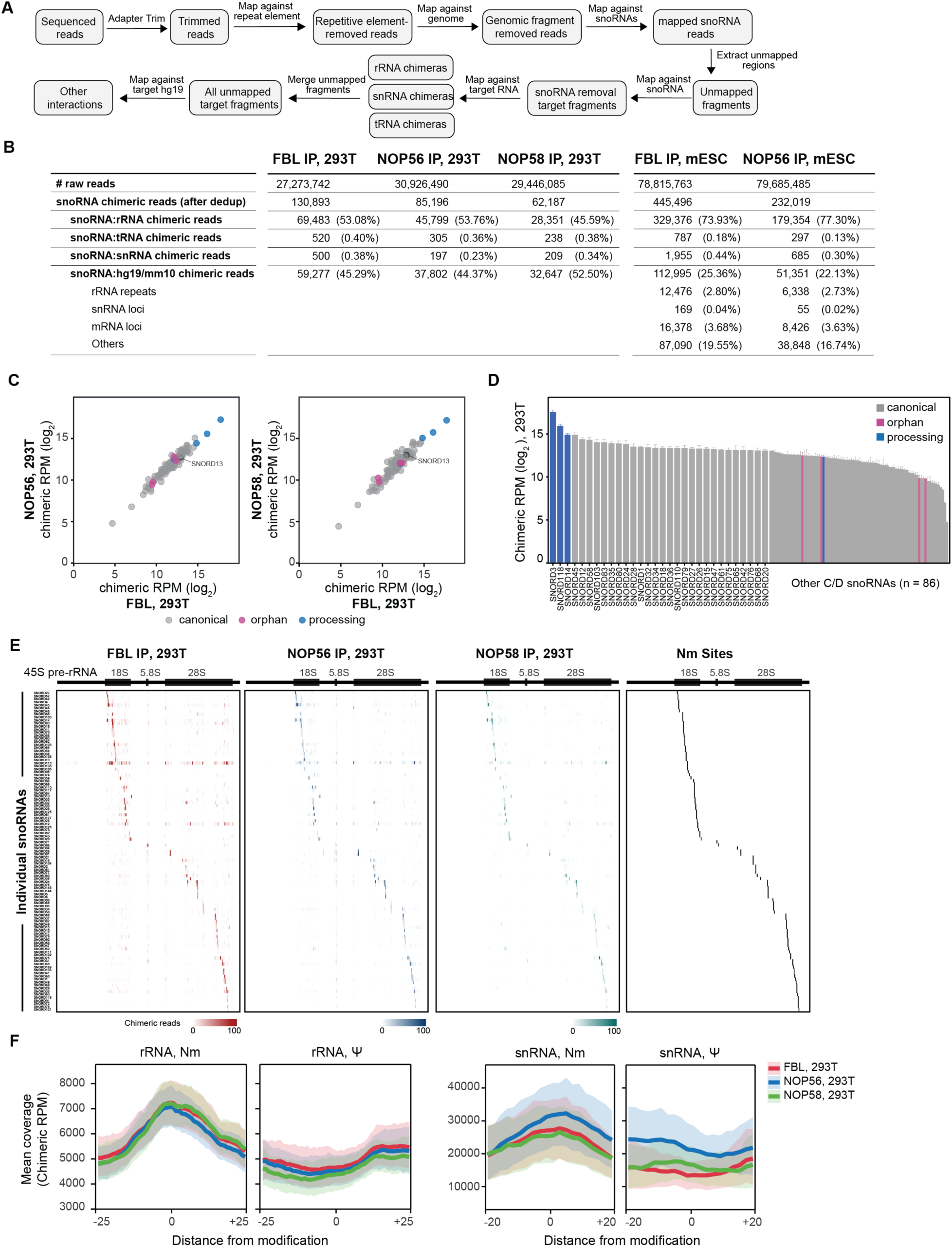
Chimeric eCLIP of core C/D snoRNP proteins comprehensively recovers known C/D snoRNA interactions in 293T cells. **(A)** Flowchart of chimeric eCLIP sequence data analysis. **(B)** Summary table of chimeric eCLIP sequencing reads. **(C)** Correlation of snoRNA chimeric read abundance from FBL, NOP56, and NOP58 chimeric eCLIP. **(D)** Chimeric read abundance for individual snoRNAs colored according to snoRNA functional class. The y-axis indicates average of chimeric RPM in FBL, NOP56, and NOP58 chimeric eCLIP. **(E)** Heatmap of chimeric read coverage across pre-rRNA from FBL, NOP56, and NOP58 chimeric eCLIP for canonical snoRNAs. Known Nm target sites are shown in black. **(F)** Metagene plots of chimeric read coverage flanking Nm and pseudouridine sites in rRNA and snRNA. RPM, reads per million.

**Figure S3.**
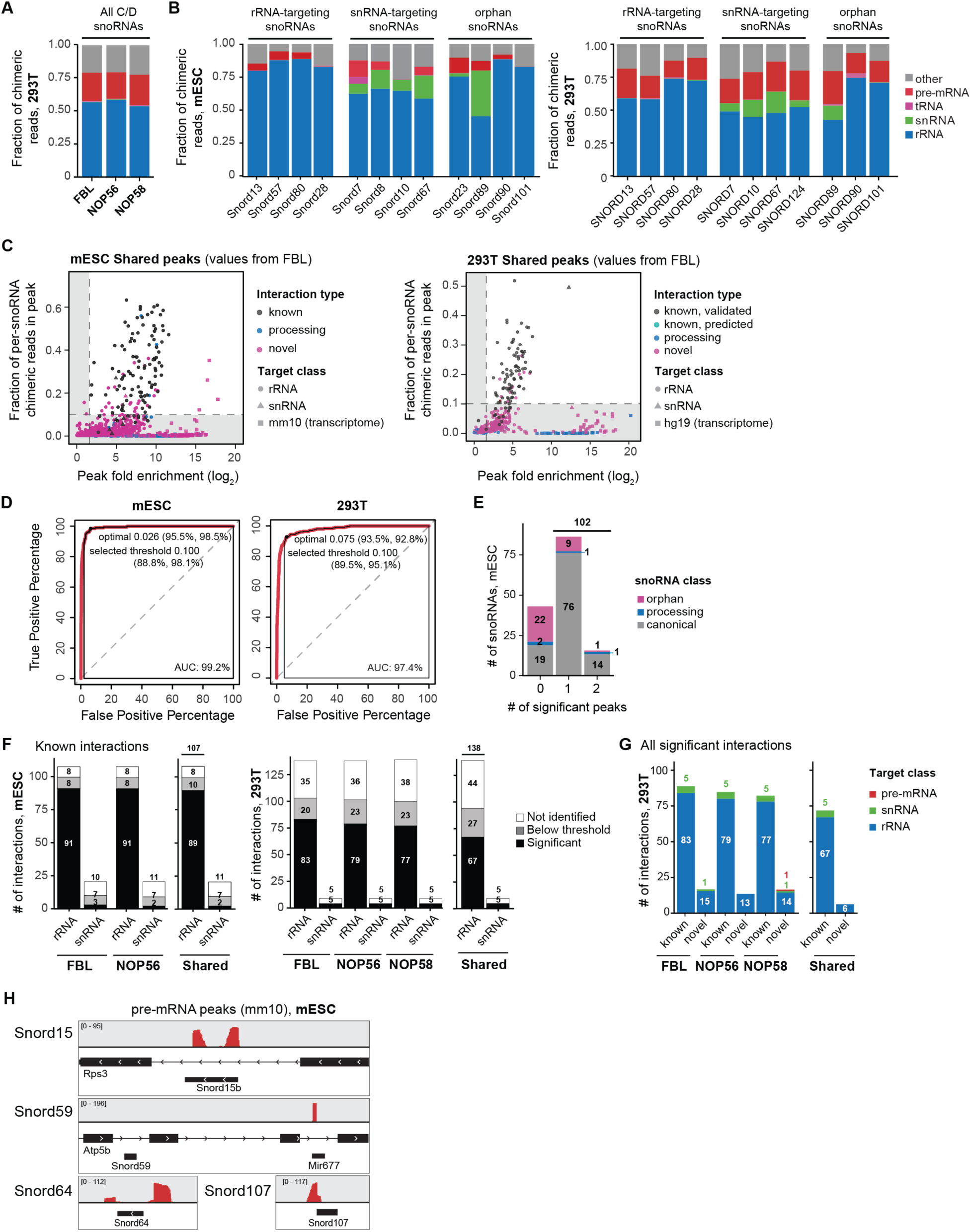
Transcriptomic mapping of C/D snoRNA chimeras in mouse and human cells. **(A)** Fraction of C/D snoRNA chimeric reads mapping to distinct classes of possible target RNAs in 293T cells from FBL, NOP56, and NOP58 chimeric eCLIP. **(B)** Distribution of individual snoRNA chimeric reads captured by FBL to distinct classes of possible target RNAs from (left) mESC and (right) 293T cells. **(C)** Implementation of “peak fold-enrichment” and “fraction of per-snoRNA chimeric reads in peak” to determine high-confidence peaks in (left) mESC and (right) 293T cells. Each datapoint indicates a peak. Colors and shapes indicate interaction status (known, previously assigned; processing, interaction of a processing snoRNA; novel, previously unassigned) and target RNA class. “Shared peaks” plots show values from FBL chimeric eCLIP of peaks confidently detected by all snoRBPs. **(D)** ROC curve to determine the threshold of the “fraction of per-snoRNA chimeric reads in peak” parameter from (left) mESC and (right) 293T cells. Chimeric eCLIP peaks overlapping known rRNA modification sites were considered as positive values to construct the curve. Optimal and selected threshold values are shown, with specificity (%) and sensitivity (%) values indicated in parentheses. **(E)** Bar chart showing the number of significant peaks per snoRNA identified in both FBL and NOP56 chimeric eCLIP from mESC. **(F)** Number of known snoRNA-target interactions identified by chimeric eCLIP for indicated snoRBP from (left) mESC and (right) 293T cells. Colors indicate status of each peak (Not identified - known interactions from database but not detected in this study; Below threshold - known interactions supported by chimeric reads but falling below threshold; and significant – known interactions overlapping high-confidence peaks above threshold established in this study). **(G)** Number of high-confidence snoRNA-target interactions identified by chimeric eCLIP in 293T cells according to interaction status and target class. **(H)** Genome browser tracks showing above-threshold putative snoRNA:pre-mRNA interactions identified in mESCs that may reflect annotation artifacts.

**Figure S4.**
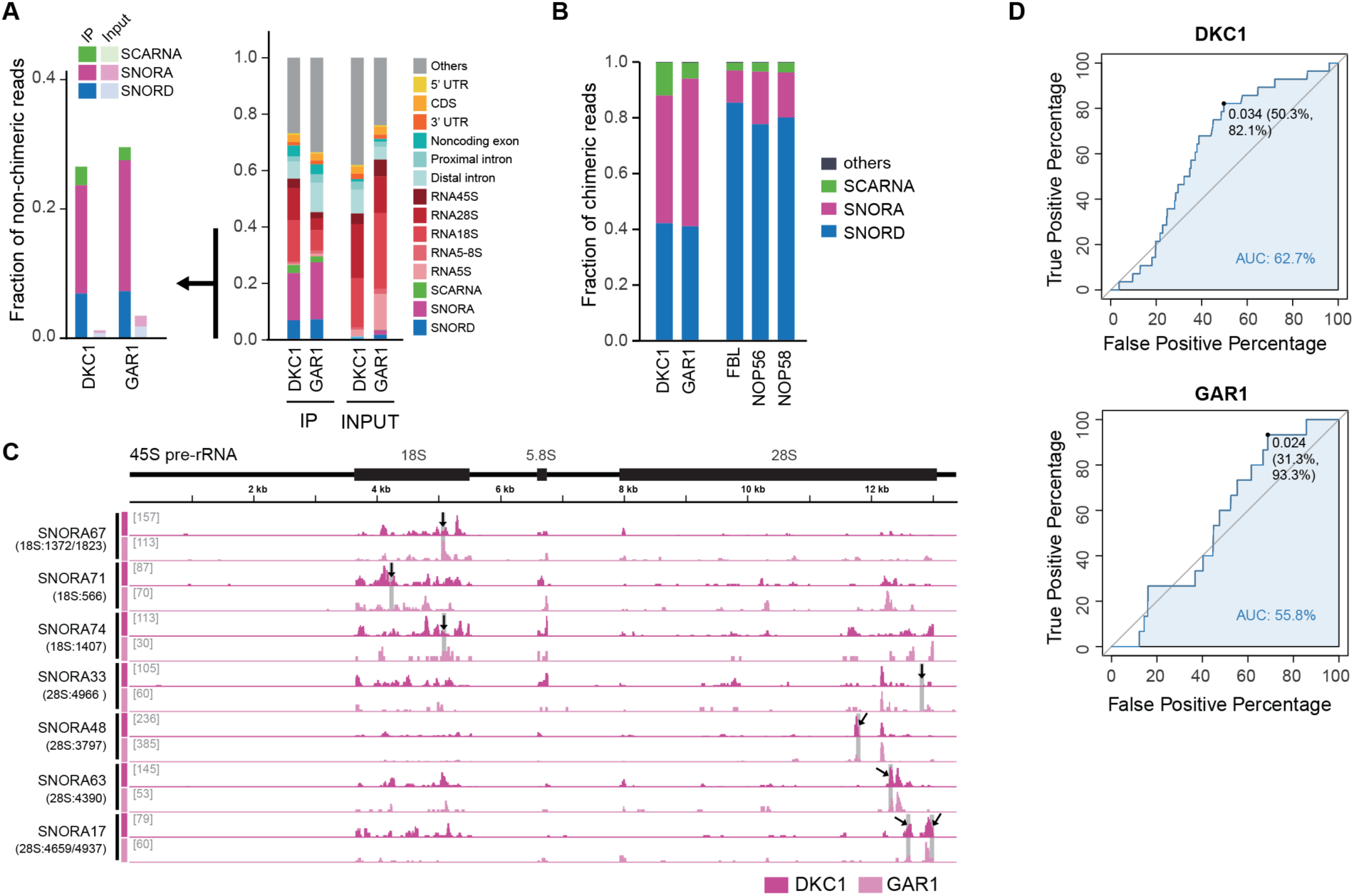
Chimeric eCLIP of core H/ACA snoRNP proteins in 293T cells. **(A)** Distribution of non-chimeric reads of (left) snoRNA classes and (right) other major RNA classes captured by DKC1 or GAR1 chimeric eCLIP. **(B)** Proportion of chimeric reads containing the indicated snoRNA classes. **(C)** Browser tracks of snoRNA chimeric reads mapped to pre-rRNA as captured by DKC1 or GAR1 chimeric eCLIP. Known pseudouridylation sites assigned to each snoRNA guide are indicated with black arrows and grey rectangles. **(D)** ROC curve from (upper) DKC1 and (bottom) GAR1 rRNA chimeric reads to determine the threshold of the “fraction of per-snoRNA chimeric reads in peak” parameter. Chimeric eCLIP peaks overlapping known rRNA modification sites were considered as positive values to construct the curve. Optimal threshold values are shown, with specificity (%) and sensitivity (%) values indicated in parentheses.

**Figure S5.**
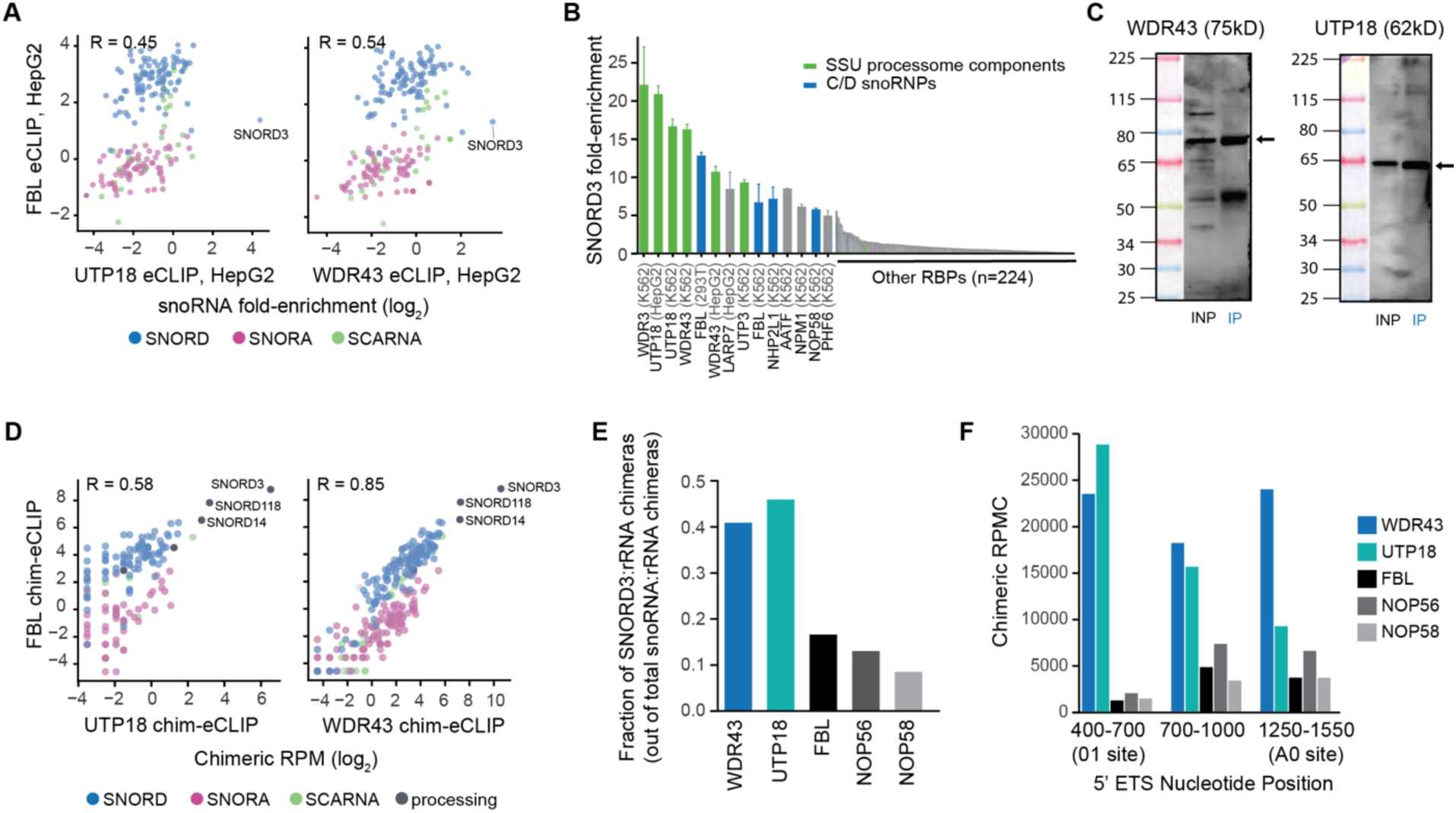
RBPs that function in ribosomal RNA processing highly enrich for SNORD3. **(A)** Correlation of snoRNA fold-enrichment in FBL versus ribosomal RNA-associated RBP in HepG2 cells. Pearson correlation (R) is shown. Only snoRNAs with 5 or more reads in IP were plotted. **(B)** Bars indicate SNORD3 fold-enrichment in eCLIP of (blue) C/D snoRNP proteins, (green) Small Subunit Processome (SSU) RBPs, and (grey) all other RBPs profiled by ENCODE. **(C)** Immunoprecipitation (IP)-western blot performed during eCLIP for indicated SSU proteins. INP, input. **(D)** Correlation of snoRNA chimeric read abundance in chimeric eCLIP of FBL versus (left) UTP18 or (right) WDR43 chimeric eCLIP in K562 cells. Pearson correlation (R) is shown. **(E)** Bars indicate the fraction of rRNA chimeric reads containing SNORD3 versus other snoRNAs captured by chimeric eCLIP using the indicated RBPs as bait. **(F)** Bars indicate the abundance of snoRNA chimeras (shown as Reads Per Million Chimeras) on different processing sites in the 5’ ETS region of pre-rRNA across the indicated RBPs.

**Figure S6.**
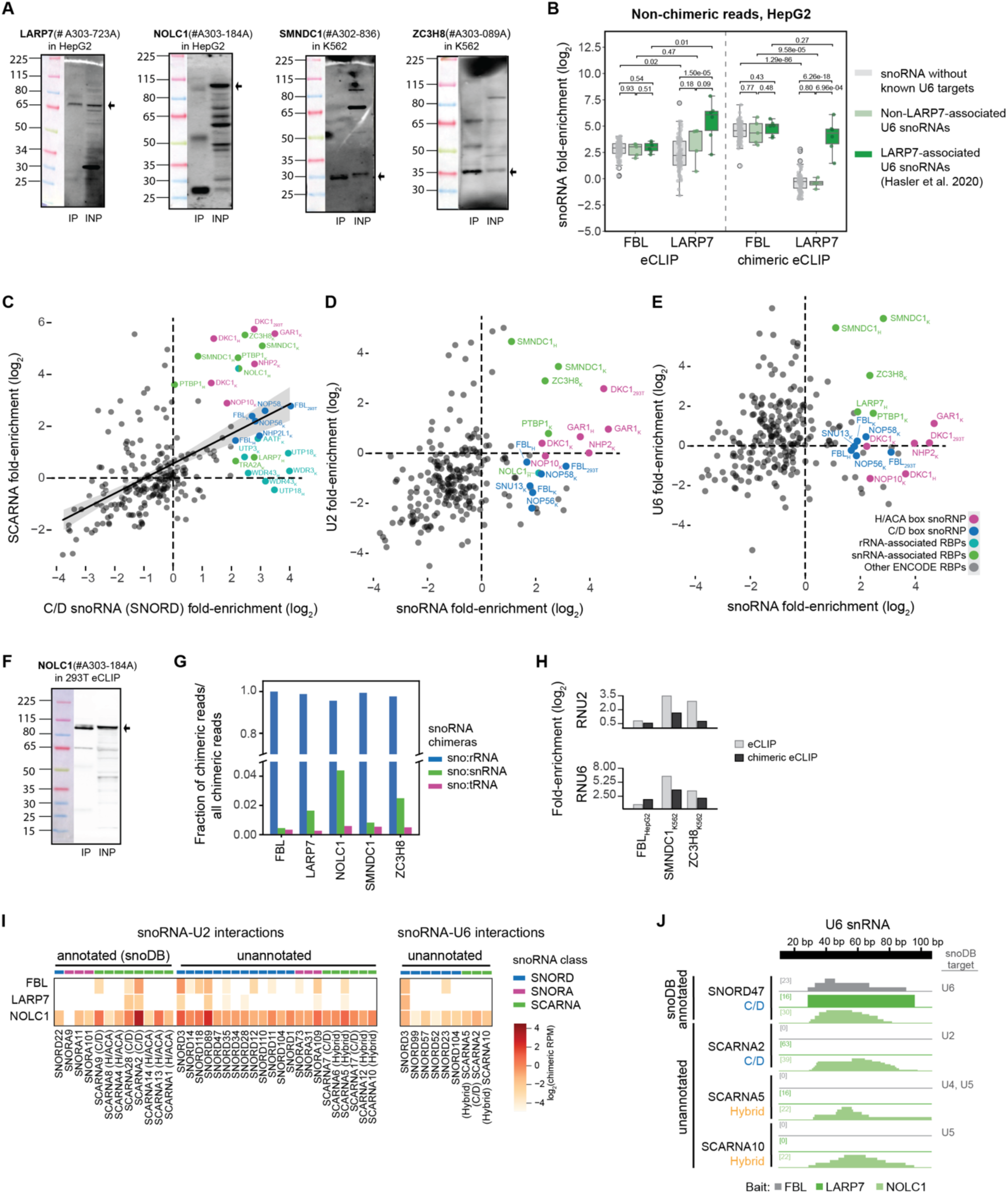
Visualization of snoRNA:snRNA chimeric reads recovered by snRNA-associated RBPs. **(A)** Immunoprecipitation (IP)-western blot performed during chimeric eCLIP for indicated proteins. INP, input. **(B)** Points indicate snoRNA fold-enrichment in (left) standard eCLIP and (right) non-chimeric reads from chimeric eCLIP recovered from FBL and LARP7 experiments in HepG2. Colors indicate snoRNAs classified as (green) LARP7-driven U6 targeting, (light green) LAPR7-independent U6 targeting, and (grey) all other snoRNAs. P-values shown are from unpaired two sample t-test. **(C)** Points indicate fold-enrichment for (x-axis) all C/D box snoRNAs versus (y-axis) SCARNAs for ENCODE eCLIP and snoRNP protein eCLIP datasets generated in this work. Labeled RBPs are as in Fig. 5A. **(D)** Points indicate fold-enrichment for (x-axis) all snoRNAs versus (y-axis) U2 snRNA for ENCODE eCLIP and snoRNP protein eCLIP datasets generated in this work. Labeled RBPs are as in Fig. 5A. **(E)** Points indicate fold-enrichment for (x-axis) all snoRNAs versus (y-axis) U6 snRNA for ENCODE eCLIP and snoRNP protein eCLIP datasets generated in this work. Labeled RBPs are as in Fig. 5A. **(F)** Immunoprecipitation (IP)-western blot performed during eCLIP for NOLC1. INP, input. **(G)** Bar indicates the fraction of chimeric reads captured by chimeric eCLIP of the indicated bait proteins that map to different target RNA classes. **(H)** Bars indicate fold-enrichment of U2 or U6 snRNA in (grey) standard eCLIP and (black) non-chimeric reads from chimeric eCLIP profiling of indicated RBPs. **(I)** Heatmap color scale indicates the read density per million sequenced reads (RPM) for (left) snoRNA:U2 chimeric reads and (right) snoRNA:U6 chimeric reads recovered by chimeric eCLIP for indicated RBPs. Cutoff for unannotated interactions shown in heatmap is 10 reads for U2 interactions and 5 reads for U6 interactions in at least one sample. **(J)** Browser tracks show read density (per million chimeric reads) for snoRNA:U6 chimeras along the U6 snRNA for known and novel candidate U6-interacting snoRNAs.

**Figure S7.**
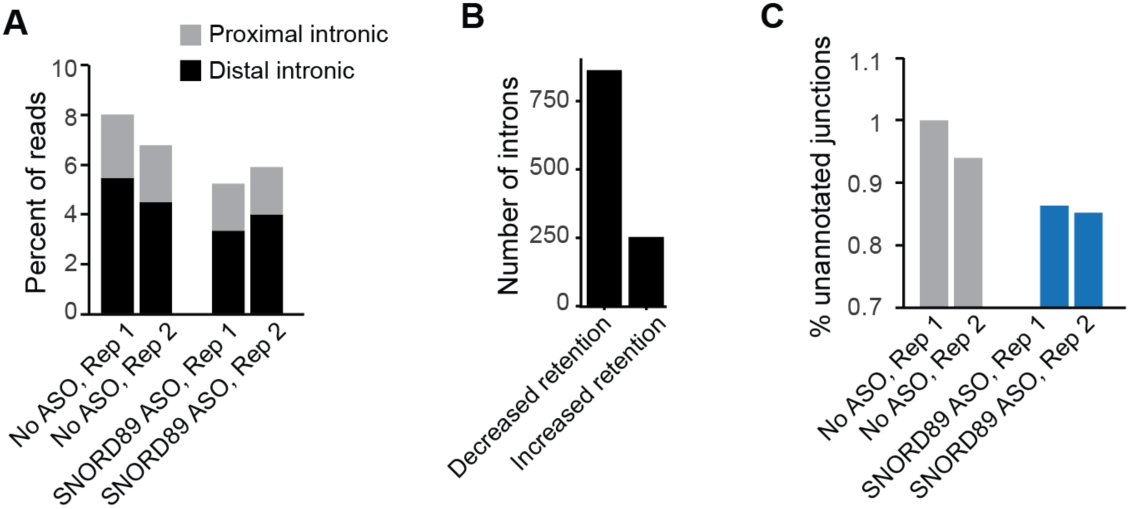
SNORD89 knockdown decreases retained introns and utilization of unannotated splice junctions. **(A)** Bars indicate the percent of mapped reads mapping to proximal (<500nt from splice site) or distal (other) intronic regions from RNA-seq performed in SNORD89 ASO or no ASO control. **(B)** Bars indicate the number of introns with significantly altered retention upon SNORD89 knockdown. **(C)** Bars indicate the percent of spliced reads for which the junction was not annotated in GENCODE (v19) or lncipedia (v5.0).

## Materials and Methods

### Cell culture

293T (Clontech), K562 (ATCC) and HepG2 (ATCC) cell lines were purchased from commercial suppliers. 293T cells were cultured in DMEM (Thermo Fisher Scientific) supplemented with 10% FBS (Cytivia). Mouse ESC lines used in this study were derived from an *M. musculus* (129/Sv) × *M. castaneous* cross and have integrated EF1α-rtTA and TetO-Ngn2 constructs. For routine passaging, mESCs were maintained on plates pre-coated with 0.2% gelatin in DMEM (Thermo) supplemented with 10 mM HEPES (Thermo), 0.11 mM β-mercaptoethanol (Sigma), 1X nonessential amino acids (Corning), 2 mM L-glutamine (Corning), 15% fetal bovine serum (Cytivia), and 1000 U/mL leukemia inhibitory factor (LIF; Sigma). All cell lines were maintained in a humidified 5% CO2 incubator at 37°C, and cell lines were crosslinked as previously described [30].

### eCLIP library preparation

eCLIP was performed as previously described [61] using antibodies against FBL (A303-891A, Fortis Life Sciences), DKC1 (GTX109000, GeneTex), NOP56 (A302-721A, Fortis Life Sciences), NOP58 (A302-718A, Fortis Life Sciences), SNU13 (A304-030A, Fortis Life Sciences), NHP2 (15128-1-AP, Proteintech), NOP10 (ab134902, Abcam), and GAR1 (11711-1-AP, Proteintech) in K562, 293T, or HepG2 cells as indicated. For all above proteins, immunoprecipitation and western blot were performed with the same antibody. Samples were sequenced on the Illumina Novaseq X Plus platform.

### eCLIP analysis

eCLIP data was analyzed using a previously described pipeline, including adapter trimming (cutadapt v3.4), removal of reads mapping to repetitive elements (STAR_2.4.0j), and mapping (STAR_2.4.0j) to the human genome (hg19) [61]. Quantitation of reads mapping to snoRNA and other repeat elements was performed according to a previously described analysis pipeline [37], using an updated database of snoRNA annotations generated by obtaining snoDB v1 [62] and removing duplicates. To compare against other RBPs, the updated snoRNA annotations were used to re-analyze all 223 eCLIP datasets (150 RBPs) generated by the ENCODE consortium [29]. snoRNA fold enrichment was calculated as reads per million in IP versus matched INPUT. For analyses comparing individual snoRNA fold-enrichment, snoRNAs with less than 5 reads in the IP sample were not included. RBP functional annotations were taken from the ENCODE analysis [37], and rRNA processing annotations were obtained from previous studies [63]. Fraction of RNA reads and enrichments shown in Fig. 1 and Fig. S1 were obtained by averaging the reads per million across two replicates. Relative information content of each element in each replicate was calculated as 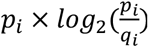, where *p_i_* and *q_i_* are the fraction of total reads in IP and input respectively that map to element *i*.

### Chimeric eCLIP library preparation

The chimeric eCLIP method is based off of the previously described miRNA chimeric eCLIP protocol [32] with modifications as described below (see Supplemental Protocol 1 for full step-by-step protocol). Briefly, 3 million cells were crosslinked with UV (254 nm, 400 mJ/cm^2^) to stabilize RNA binding protein (RBP)-RNA interactions, followed by lysis, sonication and limited digestion with RNase I (Ambion). RBP-RNA complexes were immunoprecipitated overnight with a primary antibody of interest (5 ug) precoupled to magnetic secondary antibody beads (typically 200 ul M-280 Sheep Anti-Rabbit IgG Dynabeads, Thermo). Antibodies used were mostly identical to those used for eCLIP, including: FBL (A303-891A, Fortis Life Sciences), DKC1 (GTX109000, GeneTex), NOP56 (A302-721A human and A302-720A mouse, Fortis Life Sciences), NOP58 (A302-718A, Fortis Life Sciences), GAR1 (11711-1-AP, Proteintech), ZC3H8 (A303-089A, Fortis Life Sciences), NOLC1 (A303-184A, Fortis Life Sciences), and LARP7 (A303-723A, Fortis Life Sciences); SMNDC1 used a different antibody (A302-836, Fortis Life Sciences) as the antibody used for eCLIP by the ENCODE consortium is no longer commercially available. 2% of sample was saved as for paired input control, with the remainder used for immunoprecipitation. After washes with low- and high-salt buffer, the 3’ end of RNA fragments was dephosphorylated with FastAP (Thermo) and T4 PNK (NEB), followed by 5’ end phosphorylation of cleaved target RNAs with T4 polynucleotide kinase (PNK, 3’ -phosphatase minus, NEB)). Chimeric ligation was then performed on beads at room temperature for 1 hour with T4 RNA ligase (NEB) without adapters. After additional dephosphorylation steps (FastAP, T4 PNK), phosphorylated RNA adapter was ligated to the 3’ end (T4 RNA Ligase, NEB). Samples were then denatured with 1X NuPage buffer (Life Technologies) and DTT, run on 4%–12% Bis-Tris protein gels and transferred to nitrocellulose membranes(GE Amersham).The region from the observed protein size to 75kD above (corresponding to ∼250 nt RNA) was then isolated from membrane by razor blade, and RNA was isolated by Proteinase K (NEB) treatment followed by column purification (Zymo). RNA was then reverse transcribed with SuperScript IV Reverse Transcriptase (Invitrogen) using manganese chloride-based buffer to enhance read-through of modified nucleotides [64]. Excess primers were removed with ExoSAP-IT (Thermo), and RNA was degraded by alkaline hydrolysis, followed by RNA purification (MyOne Silane beads, Thermo). Next, a 5’ Illumina DNA adapter was ligated to the 3’ end of cDNA fragment with T4 RNA ligase HC enzyme on beads overnight. After beads cleanup, an aliquot of each sample was proceeded for qPCR to identify the proper PCR cycles. The remainder of the samples were amplified with barcoded Illumina primers (Q5, NEB) and size selected via AMPure XP beads and 2% agarose gel (195-350bp). Samples were then quantified by Agilent TapeStation and sequenced either on the Illumina NextSeq (150 bp SE), NovaSeq X Plus (150 bp PE) or NovaSeq 6000 (75 bp SE) platforms.

Immunoprecipitation efficiency was tested by western blot using indicated protein antibodies at 0.2-0.5 ug/ml and Mouse or Rabbit TrueBlot secondary antibody at 1:4000.

### Chimeric eCLIP mapping

The pipeline for analysis of chimeric eCLIP datasets is available at: https://github.com/VanNostrandLab/snoRNA-chimeric, and the step-by-step pipeline is shown in Fig. S2A. Briefly, after demultiplexing using umi_tools (v1.1.1) to remove unique in-line barcode (10 random nucleotides), which was saved to read name for late use, adapter trimming was performed by cutadapt (v3.2). Next, the trimmed reads were mapped against human elements in RepBase with STAR (v2.4.0j), the reads unmapped to RepBase were then proceeded for the human genome (hg19) mapping with STAR (v2.4.0j). To remove the nonchimeric reads as much as possible, the stringent EndToEnd genomic alignment was performed and the chimeric reads with different region fragments ligated could not be aligned to genome. Those unmapped reads were then first mapped to snoRNA sequences compiled from human snoDB (v1) or mouse in-house annotations (Table S2) using bowtie (v1.3.0). The aligned snoRNA sequences were then filtered. The fragments aligned to snoRNAs over 18 nucleotides would be kept and if one read have multiple alignment results, only the reads with the maximum alignment score and longest length will be saved. In each of these reads, the remaining fragment that could not be aligned to snoRNA and over 18 nucleotides were saved as potential target regions. The potential target regions were then mapped against snoRNA sequences again to remove snoRNA: snoRNA interactions. The remaining reads were mapped to target RNA sequence (ribosomal RNA, small nuclear RNA, and tRNA) separately using bowtie2. Only the fragments mapped to target RNAs longer than 16 nucleotides and the gap between snoRNA aligned sequence and target RNA aligned sequence less than 4 were considered as real chimeric reads. Lastly, if the target fragments could not be aligned to any reference sequence, they would be merged and mapped to human (hg19) or mouse (mm10) using STAR. PCR duplicities were removed by comparing in-line barcode, sequence, snoRNA mapping location and target RNA mapping location. Unless otherwise indicated, snoRNA enrichments from FBL chimeric eCLIP in HepG2 shown were obtained by averaging the reads per million across two replicates.

### Chimeric eCLIP peak analysis

Peak calling was performed on individual snoRNA:target mapped files using Clipper [61]. To improve peak calling on edges of transcripts, all mapped chimeric rRNA/snRNA/tRNA read files, as well as the coordinates in the CLIPper reference files were “padded” (i.e. each start/end coordinate was extended by 30 nt) in silico. Peak calling on the genome was proceeded without padding. Immediately adjacent Clipper peaks were merged and considered as a single peak. Peak normalization was then performed using ‘overlap_peakfi_with_bam.pl’ script from the seCLIP pipeline [61]. Sequencing files from the input sample were processed according to the standard eCLIP pipeline and used as a background model for peak normalization. The input files were also padded prior to normalization, as processed with the IP files. Upon peak calling, the rRNA/snRNA/tRNA peak files were than “depadded” to the original positions. Peaks mapped to the genome were further subjected to filtering to discard any peaks corresponding to rRNA, snoRNA, or snRNA repeats that have escape the genome masking process.

The ‘known’ human snoRNA:target interactions were assigned based on snoDB. The mouse interactions were assigned based on previous literature[42,65–69] and is summarized in Table S2. High-confidence peaks were determined using parameters such as peak fold enrichment in the IP relative to the input (peak fc > 3) and fraction of reads at each peak for a given snoRNA (peak fraction > 0.1). An ROC was constructed using rRNA peaks to test whether the ‘peak fraction’ criteria are suitable for classifying previously known positive interactions. The peak fraction threshold was selected to minimize the false positive rate.

### Prediction of D/D’ box position and base pairing region

For D box prediction, regular expression was used to match the last occurrence of the exact CTGA consensus motif for each snoRNA in the annotation. For D’ box prediction, multi-tier regular expression was used to match all the occurrence of variations of D box motifs observed in annotated snoRNAs (i.e. CTGA, ATGA, GGTG, GTGG, GTGA, CTTA, CCGA, TCGA, CAGA, TGCA, AGGA, and TGTA) and putative antisense element was extracted 20 nt upstream of each potential D/D’ box. In the rare cases where no motif was found, regions downstream of the C box motif (GATGA) was used to assess base complementarity with the chimeric target peak. The base pairing region was determined using the in silico tool, RNAplex, to assess the complementarity between the sequence within the significant peaks and putative antisense elements. The antisense element that showed base pairing close to the D/D’ box, with low energy and high complementary proximal to the D/D’ box was selected as the putative antisense element.

### Antisense oligonucleotide treatment of mESCs

For ASO experiments, mESC were first subject to *in vitro* neuron differentiated followed by free uptake of ASO, as follows. mESC were cultured for 24 hours on gelatin-coated plates in neuron differentiation media [BrainPhys neuronal medium (Stem Cell Technologies) supplemented with 1X NeuroCult SM1 (Stem Cell Technologies), 1X N2 supplement (Thermo), 20 ng/mL brain-derived neurotrophic factor (Stem Cell Technologies)] containing 1 μg/mL doxycycline (Sigma) to induce Ngn2 expression. Then, cells were re-plated onto 0.1% PEI coated dishes in neuron differentiation media containing 1 ug/mL doxycycline (Sigma). Within 24 hours of re-plating, 5-10-5 MOE gapmer ASOs (IDT) were added to the culture media at a final concentration of 10 uM to allow for free uptake by the neurons. RNA was extracted 72 to 96 hours after addition of the ASO using TRIzol (Thermo). The ASO sequences were Snord101 ASO 5’-GGGTATCCGACAATTCAAGT-3’ and Snord89 ASO 5’-TGACTCGCTTCAGGAGGTTT-3’.

### Antisense oligonucleotide treatment of 293T cells

293T cells were transfected with 70 nM of 5-10-5 MOE gapmer ASO (IDT) using Lipofectamine RNAiMAX Transfection Reagent (Thermo). Cell culture media was replaced 24 hours after transfection, and RNA was isolated 48 hours after transfection using TRIzol (Thermo). The ASO sequence was SNORD89 ASO: 5’-TCGCTTCAGGATATTTTGTC-3’.

### RT-qPCR

RT-qPCR reactions of 12.5 ng DNase-treated RNA were prepared using the iTaq™ Universal SYBR® Green One-Step Kit (Bio-Rad) and run on the Bio-Rad CFX Opus 384 Real-Time PCR System. Relative expression was calculated using the delta-delta Ct method relative to U6 snRNA (for mESC experiments) or 5S rRNA (for 293T experiments). The primers were Snord101 (ms) 5’-GTTTGCATGATGACTTGAATTG -3’ / 5’-GTTTGTCAGACTAGCTGTTTC-3’, Snord89 (ms) 5’-CGAATTGCAGTGTCTCCATC-3’ / 5’-TGACTCGCTTCAGGAGGTTT-3’, U6 (ms) 5’-CTTCGGCAGCACATATACTAAAA-3’ / 5’-ATTTGCGTGTCATCCTTGCG-3’, SNORD89 (hu) 5’-AAAAGGCCGAATTGCAGTGT-3’ / 5’-GGTCAGACTAGTGGTTCGCT-3’, and 5S rRNA (hu) 5’-GCCATACCACCCTGAACG-3’ / 5’-AGCCTACAGCACCCGGTATT-3’.

### RiboMeth-seq (RMS)

RMS library preparation and bioinformatic analyses were conducted as described previously [34,70]. 150 to 250 ng of DNase-treated RNA was hydrolyzed in 50 mM sodium bicarbonate buffer (pH 9.2) for 1 to 12 minutes. The hydrolysis reaction was quenched in 1 mL EtOH containing 10 uL 3 M NaOAc, pH 5.2 (Thermo) and 15 ug Glycoblue coprecipitant (Thermo) and submerged in liquid nitrogen. Hydrolyzed RNA was then precipitated and washed in 80% EtOH. The intended fragment size of around 50-100 bp was verified on the TapeStation 4200 system (Agilent). RNA was dephosphorylated by Antarctic Phosphatase (New England Biolabs) for 30 minutes at 37°C, and end-repaired by T4 PNK (New England Biolabs) in the presence of 1 mM ATP (New England Biolabs) for 1 hour at 37°C. End-repaired RNA was purified using RNA Clean & Concentrator-5 columns (Zymo). Illumina libraries were then prepared using the NEBNext® Multiplex Small RNA Library Prep Set for Illumina® kit (New England Biolabs) and purified on NucleoSpin® Gel and PCR Clean-up columns (Takara Bio). Illumina sequencing of the libraries was performed on HiSeq (50 bp SE) or NovaSeq (SE100 or PE100) platforms.

### RNA-seq analysis of SNORD89 knockdown

PolyA-selected RNA-seq libraries were prepared by Novogene and sequenced on the Illumina NovaSeq (150 bp PE) platform. After adapter trimming (cutadapt v3.4, reads were first mapped (STAR, v2.4.0j) against a database of repetitive elements (RepBase v18.05), with aligned reads removed. Unmapped reads were then mapped (STAR v2.4.0j) against the human genome (hg19). Alternative splicing was quantitated with rMATS-Turbo (v4.3.0) [71] for SNORD89 ASO versus no-ASO control. Unannotated junctions were defined as splice junctions identified in STAR mapping that were not present in Gencode v19 annotations (specifically, gencode.v19.chr_patch_hapl_scaff) or lncipedia (v5.0) [72]. ENCODE RBP knockdown data was obtained from the ENCODE Consortium [29].

## Availability of data and materials

The datasets supporting the conclusions of this article are available in the Gene Expression Omnibus under accession identifiers GSE277413 (RNA-seq), GSE277414 (chimeric eCLIP), GSE277416 (eCLIP), and GSE277419 (Ribo-Meth-seq).

## Acknowledgements

We thank the Bauer Core Facility at Harvard University for technical support and access to high-throughput sequencing instrumentation. We want to thank all members of the Whipple and Van Nostrand labs for valuable discussions.

## Contributions

Conceptualization and methodology, Z.S., B.B., A.J.W., E.L.V.N; Investigation and Formal Analysis, Z.S., B.B., S.S., F.Y., T.D.Z., M.A., C.L.R.; Writing-Original draft, Z.S., B.B., A.J.W., E.L.V.N.; Writing-Review and editing, A.J.W., E.L.V.N.; Supervision and funding acquisition, A.J.W., E.L.V.N.

