## Supplemental Protocol 1 for "Mapping snoRNA-target RNA interactions in an RNA binding protein-dependent manner with chimeric eCLIP"

### Chimeric eCLIP (snoRNA) Experimental Procedures

#### Buffers & Solutions

#### Enzymes

P = user-prepared personal stock

S = stock from enzyme kit

A = aliquots stored in -80 freezer

**P iCLIP Lysis Buffer**
50 mM Tris-HCl pH 7.4
100 mM NaCl
1% NP-40 (Igepal CA630)
0.1% SDS
0.5% sodium deoxycholate (protect from light)

**P High Salt Wash Buffer**
50 mM Tris-HCl pH 7.4
1 M NaCl
1 mM EDTA
1% NP-40
P 0.1% SDS
0.5% sodium deoxycholate (protect from light)

**Wash Buffer**
20 mM Tris-HCl pH 7.4
10 mM $MgCl_2$
0.2% Tween-20
5 mM NaCl

**4x Mn SSIII/IV buffer (make fresh)**
**should be prepared fresh just before use*
200 mM Tris pH 8
300 mM KCl
12 mM $MnCl_2$

**RLTW Buffer**
1x Qiagen cat # 79216
0.025% Tween-20

**Transfer Buffer**
50 mL (1:20 of 20x) NuPage Transfer Buffer
850 mL distilled $H_2O$
100 mL 100% Methanol

**P 80% Ethanol**

**P 1% Tween-20**

**P 1M HCl**

**P 1M NaOH**

**P PKS Buffer**
100mM Tris-HCl pH 7.4
50mM NaCl
10mM EDTA
0.2% SDS

**P TT Elution Buffer**
10 mM Tris pH 7.5
0.01% Tween-20
0.1 mM EDTA

| | | | |
|---|---|---|---|
| **Turbo DNase** | 2 U/µl | LifeTech | AM2239 |
| **RNase I** | 100 U/µl | LifeTech | AM2295 |
| **FastAP** | 1 U/µl | LifeTech | EF0652 |
| **Murine RNase Inhibitor** | 40 U/µl | NEB | M0314L |
| **T4 PNK** | 10 U/µl | NEB | M0201L |
| **T4 Polynucleotide Kinase (3' phosphatase minus)** | | NEB | M0236S |
| **T4 RNA ligase 1 high conc** | 30 U/µl | NEB | M0437M |
| **Proteinase K** | 0.8 U/µl | NEB | P8107S |
| **Q5 PCR Master Mix** | | NEB | M0492L |
| **Protease Inhibitor Cocktail III** | | EMD Millipore | 539134-1SET |
| **SuperScript III Reverse Transcriptase** | | LifeTech | 18080044 |
| **Exo-SAP-IT** | | Affymetrix | 78201 |

#### Beads

| | | | |
|---|---|---|---|
| Dynabeads M-280 sheep anti-rabbit | 10 mg/ml | LifeTech | 11204D |
| (or Dynabeads Protein G) | 30 mg/ml | LifeTech | |
| Dynabeads MyOne Silane | 40 mg/ml | LifeTech | 37002D |
| Agencourt AMPure XP beads | | Beckman Coųlter | A63881 |

#### Antibodies

| | | | |
|---|---|---|---|
| FBL | 1000 µg/ml | Bethyl | A303-891A |

### Primers

#### RNA oligos:

##### Original RNA adapters:

InvRiL19: /5Phos/rArGrArUrCrGrGrArArGrArGrCrArCrArCrGrUrC/3SpC3/

**(Order 100 nmole RNA oligo, standard desalting; storage stock 200 µM; final concentration 1 µM (input), 4 µM (CLIP)).**

#### DNA oligos:

InvRand3Tr3: /5Phos/NNNNNNNNNNAGATCGGAAGAGCGTCGTGT/3SpC3/

**(Order 100 nmole DNA oligo, standard desalting; storage stock 200 µM; working stock 80 µM; final concentration 3 µM).**

InvAR17: CAGACGTGTGCTCTTCCGA **(25 nmole DNA oligo, standard desalting; storage stock 200 µM; working stock 5 µM; final concentration 0.5 µM).**

**(Below we order page-purified)**

| | |
|---|---|
| PCR_F_D501 | AATGATACGGCGACCACCGAGATCTACACTATAGCCTACACTCTTTCCCTACACGACGCTCTTCCGATCT |
| PCR_F_D502 | AATGATACGGCGACCACCGAGATCTACACATAGAGGCACACTCTTTCCCTACACGACGCTCTTCCGATCT |
| PCR_R_D701 | CAAGCAGAAGACGGCATACGAGATCGAGTAATGTGACTGGAGTTCAGACGTGTGCTCTTCCGATC |
| PCR_R_D702 | CAAGCAGAAGACGGCATACGAGATTCTCCGGAGTGACTGGAGTTCAGACGTGTGCTCTTCCGATC |

**(See Illumina customer service letter for D503-508, D703-712; any standard Illumina HT RNA-seq primers work fine)**

**(stock 100 uM; working 20 uM)**

### Notes

- Although we do not standardly do P32 labeling, we still use the “HOT” and “COLD” membranes nomenclature from iCLIP & other CLIP protocols as a shortcut. COLD = an analytical gel run on 10% of sample run as a standard Western blot; HOT = a preparative gel run on 80% of sample run for membrane cutting & RNA isolation
- When working with Protein G or other antibody-attached beads, be sure beads are never heated or dried (make sure master mixes are ready prior to removing supernatant from final washes)
- Bead washes should be done in the following order:
    1. Remove sample from magnet and buffer to beads
    2. Close tubes well and invert or flick tubes (do not vortex) to mix beads with buffer until solution is homogenous
    3. Transfer tubes to magnet and magnetically separate for 1-2 minutes. During this time, it is recommended to slowly invert magnet with tubes a few times (this helps to collect any beads from tube caps).
    4. When the solution is transparent, remove most solution while ensuring the pipette tip does not touch the beads. Next, close the tubes and spin briefly for 1-3 seconds (desktop minicentrifuge), and place back on magnet. The remaining solution can then be removed using 200 µL (or smaller) pipette tips.
- Enzymes should be kept at -20C, preferentially in chilled enzyme coolers, keep them in cooler the whole time.
- We recommend the use of low-retention tips and tubes to ensure minimal sample loss
- Ensure proper hygiene for working with RNA samples (including separation from bacteria or other RNase-containing samples) as well as high-throughput sequencing libraries (we recommend physical separation of work performed on pre-amplified and post-PCR amplified material).

### DAY 0

#### Prepare iCLIP lysis mix

- Pre-chill iCLIP Lysis Buffer
- Per sample (3 million cells): add **2.25 µl 200x Protease Inhibitor Cocktail III** and **5.5 µl Murine RNase Inhibitor** to **500 uL iCLIP Lysis Buffer**, mix
    - ** Note: RNase Inhibitor may need to be further increased for particularly difficult samples (e.g. Pancreas).

#### Couple antibody to magnetic beads (do this first)

Note: Process IgG identically to antibodies

- **Beads and antibodies:**
    - Use **200 µl beads** per sample
        - rabbit antibodies: use sheep anti-rabbit Dynabeads
        - mouse antibodies: use sheep anti-mouse Dynabeads
    - Use **5 µg antibody** per sample

- **Prepare beads:**
    - Magnetically separate beads, remove supernatant
    - Wash beads 2x in 500 µl cold **iCLIP Lysis Buffer**
    - Resuspend beads in cold **iCLIP Lysis Buffer,** 100 µl per sample

- **Bind antibody:**
    - Add antibody (5 µg per sample) to washed beads
    - Rotate at room temp, 45 min

#### Lyse cells (while ab and beads are binding)

- **Lyse cells:**
    - Retrieve cell pellets from -80°C freezer, immediately add 500 mL cold **iCLIP Lysis Buffer + Protease Inhibitor + RNase Inhibitor Mix** to each pellet, pipette to resuspend

      **2 Pellets per experiment:**
        - Sample 1: IP-A (UV-crosslinked batch #1)
        - Sample 2: IP-B (UV-crosslinked batch #2)
    - Lyse 5 mins on ice

#### RNase treat lysate (while ab+bead binding):

- Sonicate in Bioruptor on 'low' setting, 4°C, 5 min, 30sec on / 30 sec off
- Dilute RNase I in PBS at 1:100** on ice; use 10 µl diluted RNase I per sample
    - ** This likely needs to be optimized – this is for 293Ts based on our internal optimization, neurons likely need way less
- To lysed sample(s), add 2.5 µl **Turbo Dnase** (mix immediately before use)
- To lysed sample(s), add 5 µl **diluted RNase I**, mix & immediately proceed to next step
- Incubate in Thermomixer at 1200 rpm, 37°C, 5 mins (exactly), place on ice
- Centrifuge 15,000g, 4°C, 3min
- Transfer cleared lysate to a new tube

#### Capture RBP-RNA complexes on beads

- Wash antibody beads 2x in 500 µl cold **iCLIP Lysis Buffer**
- Add cleared lysates to washed antibody beads
- Rotate 4°C, 2 h or overnight (in cold room)

### DAY 1

#### Step: SAVE INPUT SAMPLES

- Mix samples well
- To new tube, take 20 µL (2%) of Sample 1 (A-Input), 2 (B-Input) for 'HOT' (RNA) gel, store at 4°C
- To new tube, take 20 uL (2%) of Sample 1 (A-Input), 2 (B-Input) for 'COLD' (western) gel; store at 4°C
    - Take additional sample as needed to ensure being able to run any desired IP-WBs

##### Wash beads

- Wash 2x with 500 µL cold **High Salt Wash Buffer**
- Wash 3x with 500 µL cold **Wash Buffer**

##### FastAP treat beads (all samples except IgG)

- Prepare **FastAP reaction mix** on ice; 50 µl per sample:
    - $H_2O$ 38 µl
    - 10X FastAP buffer 5 µl
    - Murine RNase Inhibitor 2 µl
    - Turbo DNase 2 µl
    - FastAP enzyme 3 µl

- Mix, add **50 µl** to each sample, incubate in Thermomixer at 1200 rpm, 37°C, 10 min

##### PNK treat beads

- While beads are incubating, prepare **PNK master mix** on ice; 150 µl per sample:
    - $H_2O$ 126 µl
    - 10X PNK7 buffer 20 µl
    - T4 PNK enzyme 4 µl

- Mix, add **150 µl** to each sample (don't remove FastAP mix), incubate at 1200 rpm, 37°C, 20 min

##### Wash beads

- Add 200 uL cold **High Salt Wash Buffer**, mix, magnetically separate bead suspension, remove supernatant
- Wash 3x with 500 µL cold **Wash Buffer**
- Prepare the 3' ligation master mix
- Just before adding the 3' ligation master mix, briefly spin tubes in minifuge, magnetically separate, remove residual liquid with fine tip

##### T4PNK Minus rxn (on-bead)

- Prepare **T4PNKmius mix** on ice; 100 µl per sample:
    - $H_2O$ 84.6 µl
    - 10x PNK pH 7 Buffer 10 µl
    - 0.1M ATP 1 µl
    - 5M NaCl 0.4 µl
    - Murine RNase Inhibitor 1 µl
    - T4 PNK Minus (M0236S) enzyme 3 µl
- Mix, add **100 µl** to each sample, incubate in Thermomixer at 1200 rpm, 37°C, 20 min

##### Wash beads

- Wash 2x with 500 µL cold **Wash Buffer**

##### RNA chimeric ligation (on-bead)

- Prepare **RNA ligase mix** on bench; 180 µl per sample:
    - H2O 80.4 µl
    - 10x RNA Ligase Buffer 18 µl
    - 1% Tween-20 3.6 µl
    - DMSO 5.4 µl
    - 100 mM ATP 1.8 µl
    - 50% PEG-8000 54 µl
    - Murine RNase Inhibitor 2.4 µl
    - T4 RNA Ligase HC enzyme 14.4 µl
- Mix, add **180 µl** to each sample, incubate room temp in rotator for 1-2 hr.

##### Wash beads

- Wash 2x with 500 µL cold **High Salt Wash Buffer**
- Wash 3x with 500 µL cold **Wash Buffer**

##### FastAP treat beads (all samples except IgG)

- Prepare **FastAP reaction mix** on ice; 50 µl per sample:
    - $H_2O$ 38 µl
    - 10X FastAP buffer 5 µl
    - Murine RNase Inhibitor 2 µl
    - Turbo DNase 2 µl
    - FastAP enzyme 3 µl
- Mix, add **50 µl** to each sample, incubate in Thermomixer at 1200 rpm, 37°C, 10 min

##### PNK treat beads

- While beads are incubating, prepare **PNK master mix** on ice; 150 µl per sample:
    - $H_2O$ 126 µl
    - 10X PNK7 buffer 20 µl
    - T4 PNK enzyme 4 µl
- Mix, add **150 µl** to each sample (don't remove FastAP mix), incubate at 1200 rpm, 37°C, 20 min

##### Wash beads

- Add 200 uL cold **High Salt Buffer**, mix, magnetically separate bead suspension, remove supernatant
- Wash 3x with 500 µL cold **Wash Buffer**
- Prepare the 3' ligation master mix
- Just before adding the 3' ligation master mix, briefly spin tubes in minifuge, magnetically separate, remove residual liquid with fine tip

##### Ligate 3' RNA linker (on-bead)

- Prepare **3' ligation master mix** at room temperature (<u>not</u> on ice); 25 µl per sample:
    - $H_2O$ 8.4 µl
    - 10x RNA Ligase Buffer 3 µl
    - 1% Tween-20 0.6 µl
    - DMSO 0.9 µl
    - 100 mM ATP 0.3 µl
    - 50% PEG-8000 9 µl
    - Murine RNase Inhibitor 0.4 µl
    - RNA Ligase high conc. 2.4 µl
- Mix carefully by pipetting or flicking (do not vortex) and add **25 µl** to each sample
- To each sample, add

  - 0.3 µl TT elution buffer
  - 2.085 µl $H_2O$
  - 0.615 µl InvRiL19 (200 µM)
- Incubate at room temperature for 75 min; flick to mix every ~10 min or rotate constantly

##### Wash beads (resume IgG sample here)

- Wash 2x with 500 µL cold **Wash Buffer**, magnetically separate, remove supernatant
- Wash 2x with 500 µL cold **High Salt Wash Buffer**
- Wash 2x with 500 µL cold **Wash Buffer**

##### Prepare samples for gel loading

- **IP-Bead samples (HOT and COLD)**:
  **** Note: HOT & COLD are named relative to iCLIP gels; neither is radioactive in eCLIP**
  **HOT = CLIP gel** – for membrane transfer & RNA isolation
  **COLD = WESTERN gel** – for western imaging
  - **Remove s/n,** add **100** µl cold **Wash Buffer**, resuspend beads well
  - Move 20 µl to new tube #1 = **COLD IP Bead samples**
  - For **HOT IP Bead samples,** remove remaining 80 µl Wash Buffer and add 20 µl Wash Buffer

**Prepare input and IP samples for SDS-PAGE corresponding to following table:**

| **Final Sample Composition for Gel Loading** | | | | | | |
|---|---|---|---|---|---|---|
| **Buffer** | **IgG Bead *(if applicable)*** | **Cold 0.1% Input *(if applicable)*** | **Cold Inputs** | **Cold IP Beads** | **HOT Inputs** | **HOT IP Beads** |
| Wash Buffer | 100 µl | 20 µl *(add 18 µl to 2 µl)* | 20 µl | 20 µl | 20 µl | 20 µl |
| 4x NuPAGE LDS Buffer | 37.5 µl | 7.5 µl | 7.5 µl | 7.5 µl | 7.5 µl | 7.5 µl |
| 1M DTT | 15 µl | 3 µl | 3 µl | 3 µl | 3 µl | 3 µl |

- Denature all samples in Thermomixer, 1200 rpm, 70°C, 10 min
- Cool on ice 1 min, spin briefly in minifuge
- For **all samples**, magnetically separate prior to loading (IP AND Inputs have beads)

**Potential (not recommended) -20 C stopping point**

**Load and run gels** **your loading scheme may vary depending on conditions**

- **Load HOT gel (4-12% Bis-Tris, 10-well, 1.5 mm)**

| **HOT GEL** | 1 | 2 | 3 | 4 | 5 | 6 | 7 | 8 | 9 | 10 |
|---|---|---|---|---|---|---|---|---|---|---|
| Sample | M | **A -IP** | m | **B-IP** | m | **A-Input** | m | **B-input** | M | m |
| Volume to Load | 5 µl | 30 µl | 1 µl | 30 µl | 1 µl | 30 µl | 1 µl | 30 µl | 5 µl | 1 µl |
| % of Sample Represented | | 80% | | 80% | | 2% | | 2% | | |

- **Load COLD gel (4-12% Bis-Tris, 10-well, 1.5 mm)**

| **COLD GEL** | 1 | 2 | 3 | 4 | 5 | 6 | 7 | 8 | 9 | 10 |
|---|---|---|---|---|---|---|---|---|---|---|
| Sample | M | **A-IP** | **A-INPUT** | **B-IP** | **B-INPUT** | M | | | | |
| Volume to Load | 5 µl | 15 µl | 15 µl | 15 µl | 15 µl | 5 µl | | | | |
| % of Sample Represented | | 10% | 1% | 10% | 1% | | | | | |

- Save remaining 15 µl of COLD samples at -20 C for backup
- Run at 150V in 1X MOPS Running Buffer, 75 min or until dye front is at the bottom

##### Transfer to membranes

- **COLD gel: iBlot transfer (see Quick Reference guide for pictures and more details)**
  - iBlot2 'mini' stacks are for 1 gel, 'regular' stacks are for 2 gels
  - Unseal the Transfer Stack
  - Take the top stock off and place it to the side
  - Place bottom stack (in plastic tray) into the iBlot2 machine
  - Crack open the NuPAGE gel cassette and remove the comb and bottom 'bump' areas
  - Carefully wet the gel in DI water and place it on top of the bottom iBlot stack
  - Soak an iBlot Filter Paper in DI water, and place it on top of the bottom iBlot stack
  - Remove air bubbles using the roller
  - Remove the white plastic separator from the top stack and place the top stack over the filter paper
  - Remove air bubbles using the roller
  - Place an iBlot Absorbent Pad on top of the top stack (make sure the electrical contacts are aligned properly)
  - Close the lid of the iBlot
  - Run program (P0 is our typical program)
  - After completion, check that ladder has properly transferred to membrane
  - Place membrane in western membrane case with 10 mL of appropriate blocking buffer (either Azure Chemi or Fluorescent blocking buffer)
  - Incubate on nutator at room temp, 30 min
  - Probe with primary antibody: Replace blocking buffer with fresh buffer, and add 0.2-0.5 µg/ml primary antibody (Bethyl antibodies can usually be used at 1:4000, so 2 uL for 8 mL)
  - Incubate on nutator either at room temp for 1 hr or overnight at 4ºC
- **HOT gel:**
  - Have pre-prepared COLD (4 deg) transfer buffer with methanol: 1x NuPAGE transfer buffer, 10% methanol
  - **HOT gel**: Prepare Nitrocellulose membrane(s): incubate in transfer buffer for > 1 min
  - Wet sponges and Whatman papers in transfer buffer with methanol
  - Assemble transfer stacks, from bottom to top (black side of stack holder on bottom):

    1x sponge – 2x Whatman paper – gel – membrane – 2x Whatman paper – 1x sponge

    HOT gel: Nitrocellulose membrane from GE roll stack
- **Transfer:**
  - overnight 30V (preferred) OR
  - 2 hr 200 mA (if doing this, only hook up one transfer box per power supply)

#### Day 2

- Remove HOT membrane, rinse quickly once with sterile 1X PBS, wrap in Saran wrap, store at -20C

##### Develop COLD membrane

- Block in Azure blocking buffer, room temp, 30 min
- Probe with primary antibody: 0.2-0.5 µg/ml in Azure blocking buffer, room temp, 1 hr.
- Wash 3x with TBST, 5 min each
- Probe with secondary antibody: 1:4000 Rabbit or Mouse TrueBlot HRP in Azure blocking buffer, room temp, 1 – 3 h
  - (Note: if western fails or signal is low, 1:1000 gives higher signal)
- Wash 3x with TBST, 5 min each
- Mix equal volumes of ECL Reagent 1 + Reagent 2 (or 40:1 of ECL Plus Substrate A to Substrate B), add to membrane and incubate (mix/rotate) for 1-5 min. (1ml final volume per membrane)
- Develop 30 sec & 5 min, then judge signal (15 min maximum; if 15 sec is still too bright, expose two films

#### Cut HOT membrane

- Note RBP band on film with respect to protein markers
- Place HOT membrane on clean glass/metal surface
- Using a fresh razor blade, cut lane from HOT membrane from the RBP band to 75 kDa above it
- Slice membrane pieces into ~1-2 mm slices, use a fresh razor blade for each sample
- Transfer slices to Eppendorf tube and centrifuge – place tube on ice if doing many samples

#### Release RNA from membrane

- Prepare **Proteinase K SDS mix** on ice, 150 µl per sample:
  - PKS Buffer 130 µl
  - Proteinase K 20 µl
- Mix, add **150 µl** Proteinase K SDS mix to membrane slices, incubate in Thermomixer at 1200 rpm, 37 C, 20 min (make sure all membrane slices are submerged)
- Further incubate in Thermomixer at 1200 rpm, 50 C, for an additional 20 min
- Transfer all solution to a fresh 1.5 mL DNA loBind tube
- **IF DOING PROBE-CAPTURE, SAVE SAMPLE HERE**
  - Take desired portion of sample to new 1.5 mL tube and store at -80 until probe capture use
  - Replace volume taken with $H_2O$ to bring volume back to 150 uL
- Rinse membrane with 55 µl of water, and add to supernatant above (giving 200 µL total)

#### Zymo column cleanup – RNA Clean & Concentrator-5 columns (Cat R1016)

- Add 400 µl (2x volumes) RNA binding buffer, pipette mix well
- Add 700 µl (3.5x starting volume) of 100% ethanol & pipette mix well (take care to avoid spilling of sample)
- Transfer 650 µl of mixed sample to Zymo-Spin column
- Centrifuge 30 sec on benchtop minifuge or 5,000g
- Repeat column binding for sample: carefully pipette flow-through back onto column and centrifuge again, discard flow-through
- Repeat by reloading an additional 650 µl volume until all sample has been spun through column
- Add 400 µl RNA Prep Buffer, centrifuge for 30 sec, discard flow through
- Add 500 µl RNA Wash Buffer, centrifuge for 30 sec, discard flow through
- Add 500 µl RNA Wash Buffer, centrifuge for 30 sec, discard flow through
- Add 200 µl RNA Wash Buffer, centrifuge for 1 minute at 9,000g, discard flow through
- Centrifuge additional 2 mins
- Transfer column to new 1.5 mL tube (avoid getting Wash Buffer on column)
- **Elute: Add 10 µl H2O to column, let sit for 1 min, centrifuge for 30 sec at 9,000g**
- **Repeat elution in same eluate: take the flow-through and pipette it onto the column again, sit for 1 minute, and centrifuge 30 sec at 9,000g**
- **Place IP samples in -80 C until reverse transcription**

**Potential -80 C stopping point**

### Day 3

**START Inputs only →**

#### FastAP treat input RNA

- Prepare **FastAP master mix**; 11 µl per sample:
  - 10X FastAP Buffer 2 µl
  - $H_2O$ 6 µl
  - Murine RNase Inhibitor 1 µl
  - FastAP enzyme 2 µl

- Mix, add **11 µl** to samples, mix, incubate in Thermomixer at 1200 rpm, 37 C, 10 min

#### PNK treat input RNA

- Prepare **PNK master mix**; 75 µl per sample:
  - $H_2O$ 61 µl
  - 10X PNK Buffer 9 µl
  - Turbo DNase 1 µl
  - PNK enzyme 4 µl

- Mix, add **75 µl** to samples, mix, incubate in Thermomixer at 1200 rpm, 37 C, 20 min

#### Zymo column cleanup – RNA Clean & Concentrator-5 columns (Cat R1016)

- Add 200 µl (2x volumes) RNA binding buffer directly to 95 µL repaired RNA sample, pipette mix well
- Add 300 µl (3x starting volume) of 100% ethanol & pipette mix well (avoid spilling of sample)
- Transfer all sample to Zymo-Spin column, centrifuge 30 sec at 5,000g
- Repeat column binding for sample: carefully pipette flow-through back onto column and centrifuge again, discard flow-through
- Add 400 µl RNA Prep Buffer, centrifuge for 30 sec, discard flow through
- Add 500 µl RNA Wash Buffer, centrifuge for 30 sec, discard flow through
- Add 500 µl RNA Wash Buffer, centrifuge for 30 sec, discard flow through
- Add 200 µl RNA Wash Buffer, centrifuge for 1 minute at 9,000g, discard flow through
- Centrifuge additional 2 mins
- Transfer column to new 1.5 mL tube (avoid getting Wash Buffer on column)
- **Elute: Add 10 µl H2O to column, let sit for 1 min, centrifuge for 30 sec at 9,000g**
- **Repeat elution in same eluate: take the flow-through and pipette it onto the column again, sit for 1 minute, and centrifuge 30 sec at 9,000g**

**Potential -80 C stopping point**

#### 3' linker ligate input RNA

- **Anneal adapter:**
  - Take 5 µl of RNA (from above) – remainder of input is kept at -80 C as backup
  - To each sample, add:
    - 0.2 µl TT elution buffer
    - 0.8 µl $H_2O$
    - 0.8 µl DMSO
    - 0.2 µl InvRiL19 (200 µM)
  - Incubate 65 C, 2 min → place on ice >1 min

- **Prepare ligation master mix; 13.5 µl per sample** at room temperature (not on ice)**:**
  - $H_2O$ 2.8 µl
  - 50% PEG 8000 6 µl
  - 10X RNA Ligase Buffer 2 µl
  - 1% Tween20 0.4 µl
  - DMSO 0.6 µl
  - 100 mM ATP 0.2 µl
  - Murine RNase Inhibitor 0.3 µl
  - RNA Ligase high conc. 1.2 µl

- Flick/pipette mix, add **13.5 µl** to each sample, flick/pipette-mix, incubate at room temp for 60 min
- Flick to mix every ~15 min

#### Silane cleanup input RNA

- **Prepare beads:**
  - To 10 µl **MyONE Silane Beads** per sample, add 5x volume RLT (e.g. for 4 samples, use 40 µl of beads and add 200 µl of RLT)
  - Pipette mix, magnetically separate, and remove supernatant
  - Resuspend beads in 63 µl RLTW buffer (RLT + 0.025% Tween-20) **per sample (**e.g. for 4 samples, use 250 µl RLTW buffer). Mix well.

- **Bind RNA:**
  - Add 61 µl of bead/RLTW mixture above to each RNA sample, mix
  - Add 65 µl **100% EtOH** to each sample
  - Pipette mix 10 times, leave pipette tip in tube, pipette mix every ~3-5 min for 10 min

- **Wash beads:**
  - Magnetically separate and discard supernatant
  - Add 300 µl (PCR strip tubes) or 1 mL (1.5 mL tubes) **80% EtOH**, pipette resuspend
  - After 30 s, magnetically separate, remove supernatant
  - Repeat wash with 300 µl (PCR strip tubes) or 1 mL (1.5 mL tubes) **80% EtOH**
  - After 30 s, magnetically separate, remove supernatant
  - Wash 3rd time with 100 µl (PCR strip tubes) or 750 µl (1.5 mL tubes) **80% EtOH.**
  - Spin briefly in picoFuge, magnetically separate, remove residual liquid with fine tip
  - Dry beads well, i.e. until they stop "shining - no ethanol should be left on the bottom of strip. Beads are over-dry when they change color from brown to orange/rusty color.

- **Elute RNA:**
  - Resuspend in **9.5 µl Silane Elution Buffer**, let sit for 5 min

  Magnetically separate, transfer supernatants to strip tube(s) (will be ~**9 µl**)

**Potential -80 C stopping point**

---

**←END Inputs only**

---

All CLIP and INPUT samples are now synchronized.

##### Reverse transcribe RNA (ALL CLIP and INPUTS)

- **Anneal primer** in 8-well strip tubes:
    - To ~9µl of RNA, add:
        - 1 uL dNTP mix (10 mM each)
        - 0.5 uL RT primer InvAR17 (10 uM)
    - Heat 65 C for 2 min in pre-heated PCR block, place immediately on ice (do not cool down in PCR block)

- **Prepare RT master mix** on ice; 10 µl per sample:
    - 4x Mn SSIII/IV Buffer 5 µl
    - H2O 2.8 µl
    - 0.1M DTT 1 µl
    - Murine RNase Inhibitor 0.4 µl
    - Superscript IV Enzyme 0.8 µl

- Add 10 µl to each sample, mix, incubate 55 C, 20 min in pre-heated PCR block

##### Cleanup cDNA

- **ExoSAP Treatment**
    - Add **2.5µl ExoSAP-IT** to each sample, vortex, spin down
    - Incubate 37°C for 15 mins on PCR block
    - Add 1 µl **0.5M EDTA**, pipette-mix

- **RNA removal**
    - Add 3 µl of **1M NaOH**, pipette-mix
    - Incubate 70°C, 10 min on PCR block
    - Add 3 µl of **1M HCl**, pipette-mix (to fix pH)

##### Silane cleanup cDNA

- **Prepare beads:**
    - Take **5µl MyONE Silane beads** per sample and add 5x volume RLT buffer, mix well
    - Magnetically separate and remove supernatant
    - Resuspend beads in 93 µL **RLTW buffer per sample**

- **Bind cDNA:**
    - Add **90 µl beads+RLTW** to each sample
    - Add **108 µl 100% EtOH**
    - Pipette mix (10+ times), leave pipette tip in tube, pipette mix twice (every 5min) **for total incubation of 10 minutes at room temp**

- **Wash beads:**
    - Magnetically separate, remove supernatant
    - Wash 2× with 300 µl 80% EtOH (add 80% ethanol, move back and forth on magnet, magnetically separate, remove supernatant)
    - Wash 1x with 150 µl 80% EtOH (spin briefly in picoFuge, magnetically separate, remove residual liquid with fine tip)
    - Air-dry 5 min

##### 5' linker ligate cDNA (on-bead, in 10ul)

- **Add cDNA adapter mix**
    - To each sample, add:
        - 1.45 µl of TT Elution Buffer
        - 0.25 µl **Invrand10_3Tr3** adapter (200 uM)
        - 0.8 µl 100% **DMSO**
    - Heat at 70°C, 2 min, place immediately on ice for >1 min

- **Prepare ligation master mix** on bench:

| | |
|---|---|
| $H_2O$ | 1.4 µl |
| 10x NEB RNA Ligase Buffer (with DTT) | 1 µl |
| 0.1M DTT | 0.2 µl |
| 0.1M ATP | 0.1 µl |
| 1% Tween-20 | 0.2 µl |
| 50% PEG 8000 | 3.6 µl |
| RNA Ligase high conc (M0437M) | 1 µl |
| 5' Deadenylase (M0331S) | 0.3 µl |

- Flick to mix twice, spin down briefly, add **7.8 µl** to each sample: stir sample with pipette tip, then add master mix slowly with stirring; needs to be homogeneous
- Incubate at room temp overnight on rotator

#### Day 4

##### Silane cleanup linker-ligated cDNA

- To each sample **add 5 µl of Silane Elution Buffer,** making 15 µl total.

- **Prepare beads:**
    - Take **2.5 µl MyONE Silane beads** per sample, add 5x volume RLT
    - Magnetically separate and remove supernatant
    - Resuspend beads in 47 µL RLTW buffer per sample

- **Bind RNA:**
    - Add **45 µl** beads+**RLTW buffer mix** to each sample
    - Add **45 µl 100% EtOH**
    - Pipette mix, leave pipette tip in tube, pipette mix twice, for 10 min total

- **Wash beads:**
    - Magnetically separate, remove supernatant
    - Wash 2× with 300 µl 80% EtOH (add 80% ethanol, move back and forth on magnet, magnetically separate, remove supernatant)
    - Wash 1x with 150 µl 80% EtOH (spin briefly in picoFuge, magnetically separate, remove residual liquid with fine tip)
    - Air-dry 5 min

- **Elute ligated cDNA:**
    - Resuspend in 25 µl **Silane Elution Buffer**, let sit for 5 min
    - Magnetically separate, transfer **25 µl** sample to new tube

**Potential -80 C stopping point**

##### qPCR quantify cDNA

- Prepare **qPCR master mix**; 9 µl per sample:

| | | |
|---|---|---|
| PowerSybr 2x master mix | 5.0 µl | |
| $H_2O$ | 3.6 µl | |
| qPCR primer mix | 0.4 µl | (10 uM each qPCR-grade D5x/D7x mix) |

- Mix, dispense into 384-well qPCR plate, add **1 µl 1:10 diluted (in $H_2O$) cDNA**, seal, mix
- **qPCR conditions (preset protocol: 30 cycle, PowerSYBR, no melt)**
    - 95 C for 10 min
    - 95 C for 15 sec ← (30x)
    - 60 C for 1 min -> take image
    - **Cycle # for final PCR will be 3 cycles less than the Ct of the 1:10 diluted sample**

**** Note: we use the automatically calculated Ct for this; this '3 cycle less' rule may change based on your lab setup, so for the first couple CLIPs it is best to err on the side of 1 or 2 extra PCR cycles. If final libraries are > 50 nM (especially if > 100 nM), you should back off a couple cycles.**

#### PCR amplify cDNA

- Typical: Input 9 total cycles (6 + 3), CLIP 16 (6 + 10) total cycles
    - Note that 18 cycles will yield ~30-50% PCR duplicated libraries (further increasing above 18 cycles), which can be ok for RBPs with few specific targets but will be challenging for broad binders.
    - Cycle # for final PCR: 3 cycles less than the qPCR Ct of the 1:10 diluted sample

- Prepare **PCR** on ice; 40 µl total per sample:
    - Ligated cDNA 12 µl (save remainder at -80 C as backup)
    - $H_2O$ 4 ul
    - 20 µM right primer (D50x) 2 µl
    - 20 µM left primer (D70x) 2 µl
    - 2x Q5 PCR master mix 20 µl

- PCR conditions (cycle # depending on library):
    - 98 C for 30 s
    - 98 C for 15 sec -> 68 C for 30 sec -> 72 C for 40 sec (x6 cycles)
    - 98 C for 15 sec -> 72 C for 60 sec (x ? cycles)
    - 72 C 1 min
    - 4 C hold

#### SPRI cleanup library

- Resuspend **AmpureXP beads** well by vortexing
    a. (Note: beads should be incubated at room temp for 15 min prior to use)
- Add 72 µl bead suspension (do <u>not</u> separate) per 40 µl PCR reaction and pipette mix well
- Incubate at room temp for 10 min (pipette mix 2-3x during incubation)
- Magnetically separate, wash beads 3x with **80% EtOH**
- Dry beads for 5 min on magnet (do not over-dry, i.e. the pellet will 'crack')
- Resuspend in **20 µl of PCR Elution Buffer** and incubate for 5 minutes at room temperature
- Magnetically separate and transfer supernatant to new tubes

#### Gel-purify library

- **Prepare samples and run gel:**
    - Open E-gel EX 2% agarose gel, remove comb and load into E-gel system
    - Load 10 uL $H_2O$ or E-gel sample buffer into M (marker) lane – do not use this for ladder or sample (it tends to smear and run strangely)
    - Load 6 uL 50bp E-Gel ladder into lanes 1 and 10
    - Add 2 uL E-gel sample buffer to each sample and load samples into lanes 2-9
    - Run E-gel for 10-12 mins
    - Take picture using E-gel camera
- **Gel-extract library from gel:**
    - Crack open E-gel using spatula (take care to avoid tearing gel; if possible, keep gel on back faceplate so that the samples are in the same orientation as they were loaded)

- Under blue light illumination, cut gel slices **195**-350 bp and place into 15 mL conical tubes, using fresh razor blades for each sample; keep cross-contamination to minimum
    - Keep in mind: adapter-dimer (including RNA adapter) is **156** bp, so chimeras (20nt small RNA + 20nt target) require at least 40nt of fragment

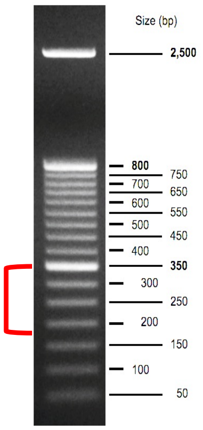


- **Cut & elute gel** using Qiagen MinElute gel extraction kit:
    - Weigh 15 mL conical with gel slice (blank with empty conical tube)
    - Calculate gel weight, add 6x volumes of **Buffer QG** to melt gel (e.g. for 100 mg gel, add 600 µl QG)
    - Melt gel at room temp (do not heat) on benchtop (can shake to help melt, but don't vortex)
    - After gel is melted, add 1x volume of original gel of **isopropanol** & mix well (100 mg gel = 100 µl isopropanol)
    - Load on column (750 µl per spin, can do multiple spins, all spins max speed 1 min)
        - **NOTE:** if gel weight is >400 mg, wash 1x with 500 µl Buffer QG after every 4 spins)
    - After all sample has been spun through, wash 1x with 500 µl **Buffer QG**
    - Add 1X with 750 µl **Buffer PE**, spin 1 min, pour out flow-through, spin again 2 min max speed
    - Carefully move column to new 1.5 mL tube (avoid any carryover of PE – if any liquid is visible on the outside of the column redo 2 min max speed spin)
    - Using a fine tip, pipette all remaining PE buffer from plastic purple rim of the MinElute column
    - Air dry 2 mins
    - Carefully add 12.5 µl **Buffer EB** directly to the center of the column, incubate 2 min room temp, spin max speed
    - For improved yield – repeat the elution (take the flow-through and add it to the column again)

#### Quantitate library (D1000 DNA tapestation)

- 3 µl D1000 loading buffer, 1 µl sample
- Vortex to mix, spin down in microfuge
- Correctly quantify by adding a region to each sample and dragging boundaries to include the entire library peak (usually ~150-600bp).

**Example of a good trace:**

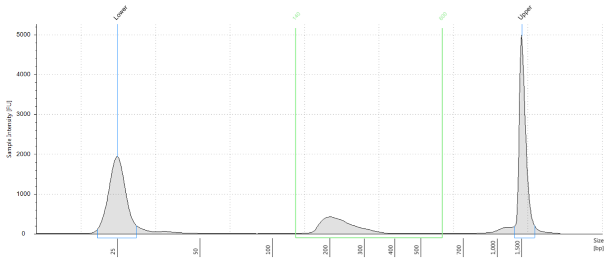


Region Table

| From [bp] | To [bp] | Average Size [bp] | Conc. [ng/µl] | Region Molarity [nmol/l] | % of Total | Region Comment | Color |
|---|---|---|---|---|---|---|---|
| 140 | 600 | 233 | 2.79 | 19.2 | 65.55 | | |
